# Cerebellar-recipient motor thalamus drives behavioral context-specific movement initiation

**DOI:** 10.1101/802124

**Authors:** Joshua Dacre, Matt Colligan, Julian Ammer, Julia Schiemann, Thomas Clarke, Victor Chamosa-Pino, Federico Claudi, J. Alex Harston, Constantinos Eleftheriou, Janelle M.P. Pakan, Cheng-Chiu Huang, Adam Hantman, Nathalie L. Rochefort, Ian Duguid

## Abstract

To initiate goal-directed behavior, animals must transform sensory cues into motor commands that generate appropriately timed actions. Sensorimotor transformations along the cerebellar-thalamocortical pathway are thought to shape motor cortical output and movement timing, but whether this pathway initiates goal-directed movement remains poorly understood. Here, we recorded and perturbed activity in cerebellar-recipient regions of motor thalamus (dentate / interpositus nucleus-recipient regions, MTh_DN/IPN_) and primary motor cortex (M1) in mice trained to execute a cued forelimb lever push task for reward. MTh_DN/IPN_ population responses were dominated by a time-locked increase in activity immediately prior to movement that was temporally uncoupled from cue presentation, providing a fixed latency feedforward motor timing signal to M1_FL_. Blocking MTh_DN/IPN_ output suppressed cued movement initiation. Stimulating the MTh_DN/IPN_ thalamocortical pathway in the absence of the cue recapitulated cue-evoked M1 membrane potential dynamics and forelimb behavior in the learned behavioral context, but generated semi-random movements in an altered behavioral context. Thus, cerebellar-recipient motor thalamocortical input to M1 is indispensable for the generation of motor commands that initiate goal-directed movement, refining our understanding of how the cerebellar-thalamocortical pathway contributes to movement timing.

## Introduction

The ability to generate precisely timed motor actions in response to sensory cues is a hallmark of mammalian motor control. Movement timing is believed to be mediated by cerebellum-dependent shaping of motor output (Holmes, 1939) given that damage to, or inactivation of, the cerebellum results in poorly timed motor actions (Bastian and Thach, 1995; Milak et al., 1997; Thach, 1975). However, the circuit mechanisms that generate motor timing signals necessary for goal-directed movement initiation remain unclear. Cerebellar control of goal-directed movement is predominantly mediated via two distinct pathways, the cerebellar-rubrospinal tract (Gibson et al., 1985) and the more dominant cerebellar-thalamocortical pathway (Horne and Butler, 1995). Feedforward excitatory input from the deep cerebellar nuclei (DCN) provides one of the main driver inputs to the motor thalamus and is thought to be necessary for controlling the timing of simple and complex movements (Ivry and Keele, 1989; Mink and Thach, 1991; Ohmae et al., 2017). But whether the cerebello-thalamocortical pathway is required for movement initiation has been much debated (Thach, 2013). Neuronal activity in the dentate and interpositus subdivisions of the deep cerebellar nuclei and their recipient regions in motor thalamus precedes activity changes in motor cortex and movement (Bosch-Bouju et al., 2014; Butler et al., 1992; Fortier et al., 1989; Harvey et al., 1979; Mushiake and Strick, 1993; Thach, 1975, 1978), suggestive of a role in movement initiation. However, local inactivation of dentate and interpositus nuclei or their recipient regions in motor thalamus during simple cued forelimb tasks produces variable behavioral outcomes, from no effect (Miller and Brooks, 1982) to slowing of reaction times (Meyer-Lohmann et al., 1977; Mink and Thach, 1991; Spidalieri et al., 1983; Thach, 1975) and reduced task engagement (van Donkelaar et al., 2000). Although suggestive of a role in movement timing, direct causal evidence supporting a role for the cerebellar-thalamocortical pathway in movement initiation has been lacking.

If the cerebellar-thalamocortical pathway conveys motor timing signals, MTh_DN/IPN_ activity could be described by three hypothetical models. Either, (i) population activity rises from a fixed timepoint prior to movement initiation which is temporally uncoupled from cue onset (i.e. a motor timing signal that has a fixed onset and fixed slope when aligned to movement); (ii) population responses rise from cue onset to a timepoint distant from movement initiation (i.e. variable onset, fixed slope trajectories where MTh_DN/IPN_ input does not directly correlate with movement initiation); or (iii) MTh_DN/IPN_ population responses change from cue onset and are directly coupled to movement initiation (i.e. linear sensorimotor transformation from cue to movement, variable onset, variable slope trajectories) (Figures 1a and 1b).

**Figure 1.**
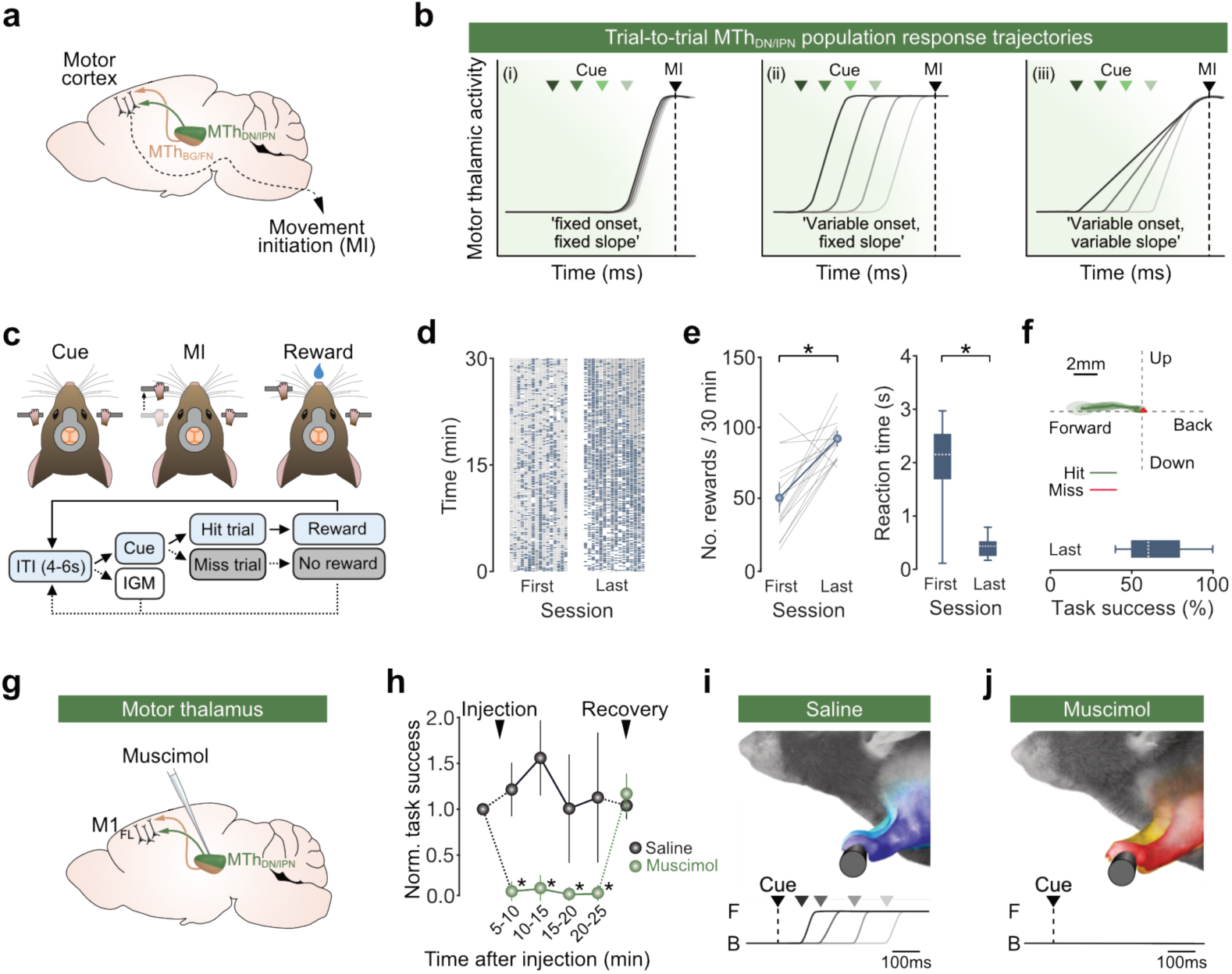
Motor thalamic output is necessary for cued goal-directed movement initiation in mice. **(a)** Sagittal mouse brain schematic depicting feedforward input from dentate / interpositus nucleus-recipient (MTh_DN/IPN_) and basal ganglia / fastigial nucleus-recipient (MTh_BG/FN_) regions of motor thalamus to motor cortex, **(b)** Hypothetical trial-to-trial MTh_DN/PN_,population response trajectories: model (i), fixed onset, fixed slope; model (ii), variable onset, fixed slope; model (iii), variable onset, variable slope. Green triangles depict cue onset across 4 trials, response trajectories are aligned to movement initiation (MI, black triangle and dashed line), **(c)** Top, cued goal-directed linear forelimb push task for head restrained mice. MI, movement initiation. Bottom, behavioral task structure: ITI, inter-trial interval; IGM, internally generated movement, **(d)** *Left & right*, rasters showing behavioral task success across first and last training sessions. Each column represents the behaviour of an individual mouse across the training session (N = 16). Blue, hit trials; grey, miss trials; white, IGMs. **(e)** Task metrics across learning. *Left*, average number of rewards received per 30 minutes (N = 16 mice, *t*(15) = -5.3,* *P* = 9.5×10^−5^. two-sample f-test). *Right*, box-and-whisker plots show median, interquartile range and range of median reaction times (RTs) across mice on the first and last day of training (N = 16 mice. *t*(15) = 7.1. * *P* = 3.4×10^−6^ two-sample f-test). **(f)** *Top*. average kinematic forepaw trajectory for hit (green) and miss (red) trials from anexample mouse shown in Supp. Video 2. Thick line depicts average trajectory overlaid with the 95% CI of frame-by-frame paw position variance (transparent ovals). *Bottom*, box-and-whisker plot showing median, interquartile range and range of task success across mice during last training session, **(g)** Focal muscimol inactivation of thalamus, targeting MTh_DN/IPN_. **(h)** Normalised task success as a function of time after muscimol injection. Colored symbols represent changes in population means ± 95% Cl after saline (black, N = 5 mice) or muscimol (green, N = 5 mice) injection (5-10 minute bin, *F*(1,8) = 63.0,’ *P* = 1.9×10^−4^,2-way repeated measures ANOVA, with a Bonferroni-Holm correction for multiple comparisons), **(i-j)** *Top:* Superimposed images of mouse forelimb position at cue onset across 3 trials after (k) saline or (I) muscimol injection targeted to MTh_DN/PN_. *Bottom:* Example lever trajectories from 4 different trials (black to light grey) following injection of (k) saline or (I) muscimol into MTh_DN/IPN_, demonstrating postural consistency despite motor thalamic inactivation. B, back lever position; F, front lever position; black triangle and dashed line represent cue presentation; grey triangles indicate reaction times.

To distinguish between these models and to investigate whether the cerebellar-recipient motor thalamocortical pathway is necessary for goal-directed movement initiation, we employed thalamic population calcium imaging, patch-clamp recordings in M1, and targeted manipulations in mice trained to execute a cued forelimb push task for reward. We demonstrate that MTh_DN/IPN_ population responses were dominated by a time-locked increase in activity immediately prior to movement initiation that was temporally uncoupled from cue presentation, providing a fixed latency feedforward motor timing signal to M1_FL_. Focal inactivation of MTh_DN/IPN_ suppressed layer 5 membrane potential dynamics in forelimb motor cortex (M1_FL_) and blocked cued movement initiation. Direct stimulation of MTh_DN/IPN_ neurons, or their axon terminals in M1_FL_, in the absence of the cue recapitulated motor cortical activity dynamics and forelimb behavior in the learned behavioral context, but generated semi-random movements in an altered behavioral context where the lever and reward were absent. Together, our findings demonstrate that the cerebellar-recipient motor thalamocortical pathway conveys essential motor timing signals necessary for the initiation of goal-directed movement.

## Results

To explore the role of MTh_DN/IPN_ in goal-directed movement initiation, we first developed a cued linear forelimb push task for mice. The design of the task, which incorporates a horizontal translation lever, required mice to learn the correct wrist and grip orientation to ensure smooth, friction-reduced horizontal lever movements (4 mm) in response to a 6 kHz auditory cue (Figure 1c and Video S1). Mice rapidly learned to execute the task (mean = 7.5 days, 95% CI [6.3, 8.6], N = 16 mice, all data, unless otherwise stated, are presented as mean, [bootstrapped 95% confidence interval]; last session task success, mean = 0.64 rewards per cue presentation, 95% CI [0.56, 0.72]), displaying relatively fast reaction times (last session, median = 0.32s [0.30, 0.34]) and reproducible forelimb kinematic trajectories (Figures 1d-1f, Video S1).

To selectively record and manipulate MTh_DN/IPN_ activity during behavior, we confirmed the anatomical location of thalamic nuclei that send monosynaptic projections to M1_FL_ and receive dense projections from the dentate (DN) and interpositus (IPN) deep cerebellar nuclei (Gao et al., 2018; Kuramoto et al., 2009; Rispal-Padel et al., 1987; Sakai et al., 1996; Schiemann et al., 2015). By employing conventional retrobead fluorescence tracing across layers 2-6 in M1_FL_, layer 5-specific monosynaptic rabies tracing and anterograde viral tracing from the DN and IPN, we observed dense expression in dorsal-posterior motor thalamus centered on the ventrolateral nucleus (VL) with expression in anteromedial (AM), ventral posteromedial (VPM) and ventral posterolateral (VPL) nuclei, but no staining in the ventromedial nucleus (VM), which primarily receives input from the basal ganglia (BG) and fastigial nucleus (FN) (Figure 1a, Figures S1 and S2) (Kuramoto et al., 2009; Person et al., 1986; Sakai et al., 1996; Tanaka et al., 2018). Moreover, the vast majority of neurons in MTh_DN/IPN_ sent direct projections to M1_FL_ highlighting the high degree of connectivity between two important nodes along the cerebellar-thalamocortical pathway (mean = 76.0%, 95% CI [69.4, 82.8], n = 16 slices from N = 3 mice, Figure S3).

To investigate whether MTh_DN/IPN_ was necessary for cued goal-directed movement initiation, we focally injected a small bolus of the GABA_A_ receptor agonist muscimol centered on the ventrolateral nucleus, with an estimated spread of ∼500 µm from the point of injection within 10 minutes (Martin, 1991) (Figure 1g and Figure S4). By applying muscimol during behavioral task engagement, we recorded the immediate post-injection effects. Muscimol reduced task success by ∼90%, 5-10 minutes after injection, an effect that persisted for the duration of the session before reverting to baseline after 24 hours (5-10 mins, mean = 0.11 normalized task success, 95% CI [0.04, 0.18, N = 5 mice, *F*(1,8) = 63.0, *P* = 1.9×10^−4^, two-way ANOVA with Bonferroni-Holm correction for multiple comparisons) (Figure 1h). Reduced task success was not a result of task disengagement as cue presentation reproducibly evoked short-latency whisking and enhanced arousal (Video S2). Moreover, mice did not experience a loss of forelimb postural control, as evidenced by the accurate trial-to-trial forelimb positioning at cue presentation (Figures 1i and 1j, Figure S4). The predominant effect of MTh_DN/IPN_ inactivation was not a slowing of reaction times (Meyer-Lohmann et al., 1977; Mink and Thach, 1991; Spidalieri et al., 1983; Thach, 1975; van Donkelaar et al., 2000), instead it selectively blocked movement initiation. Given that MTh_DN/IPN_ provides feedforward excitation to M1_FL_, we next assessed whether motor cortical output was also necessary by focally injecting muscimol into the center of M1_FL_ (Schiemann et al., 2015). M1_FL_ inactivation reduced task success by ∼70%, 5-10 minutes after injection, persisting for the duration of the session before reverting to baseline after 24 hours (5-10 mins, mean = 0.29 normalized task success, 95% CI [0.11, 0.51], N = 5 mice, *F*(1,8) = 3.7, *P* = 0.09, two-way ANOVA with Bonferroni-Holm correction for multiple comparisons). Dissimilar to motor thalamic inactivation, silencing M1_FL_ output induced a loss of forelimb postural control and hemiplegia, resulting in an inability to engage with the lever and task (Figure S4 and Video S3). Together, these data suggest that M1 is essential for coordinated motor control but MTh_DN/IPN_ output is a prerequisite for goal-directed movement initiation in mice.

To test whether MTh_DN/IPN_ response timing was consistent with a role in movement initiation, we employed Gradient Refractive Index (GRIN) lens-mediated 2-photon population calcium imaging of MTh_DN/IPN_ neurons during task engagement (Figures 2a and 2b). We found that the majority of MTh_DN/IPN_ neurons displayed task-related activity changes (192/248 neurons, 11 fields of view (FOV), N = 8 mice) either prior to movement initiation (early onset positive ΔF/F_0_, 127/192 neurons; early onset negative ΔF/F_0_, 24/192 neurons) or during the post-movement period (late onset positive ΔF/F_0_, 18/192 neurons; late onset negative ΔF/F_0_, 23/192 neurons) (Figures 2c and 2d). The most prominent activity profile of early onset neurons – i.e. activity that could contribute to movement initiation – was enhanced activity that occurred after the cue but ∼300 ms prior to movement initiation (early onset neurons – 84.1% enhanced, 15.9% suppressed) (Figure 2d). This dominant population response profile was consistent across trials and was not spatially restricted to a defined region of MTh_DN/IPN_ (Figures 2c-2e). To investigate whether MTh_DN/IPN_ population responses conveyed information regarding the cue or a purely movement-related signal, we exploited miss trials where mice perceived the cue, as indicated by short-latency whisking and increased arousal, but did not engage in the task (see Video S1). In the absence of movement, no appreciable cue-evoked responses across early onset neurons were observed (Figures 2c and 2f), suggesting MTh_DN/IPN_ output conveys a purely motor-related feedforward signal.

**Figure 2.**
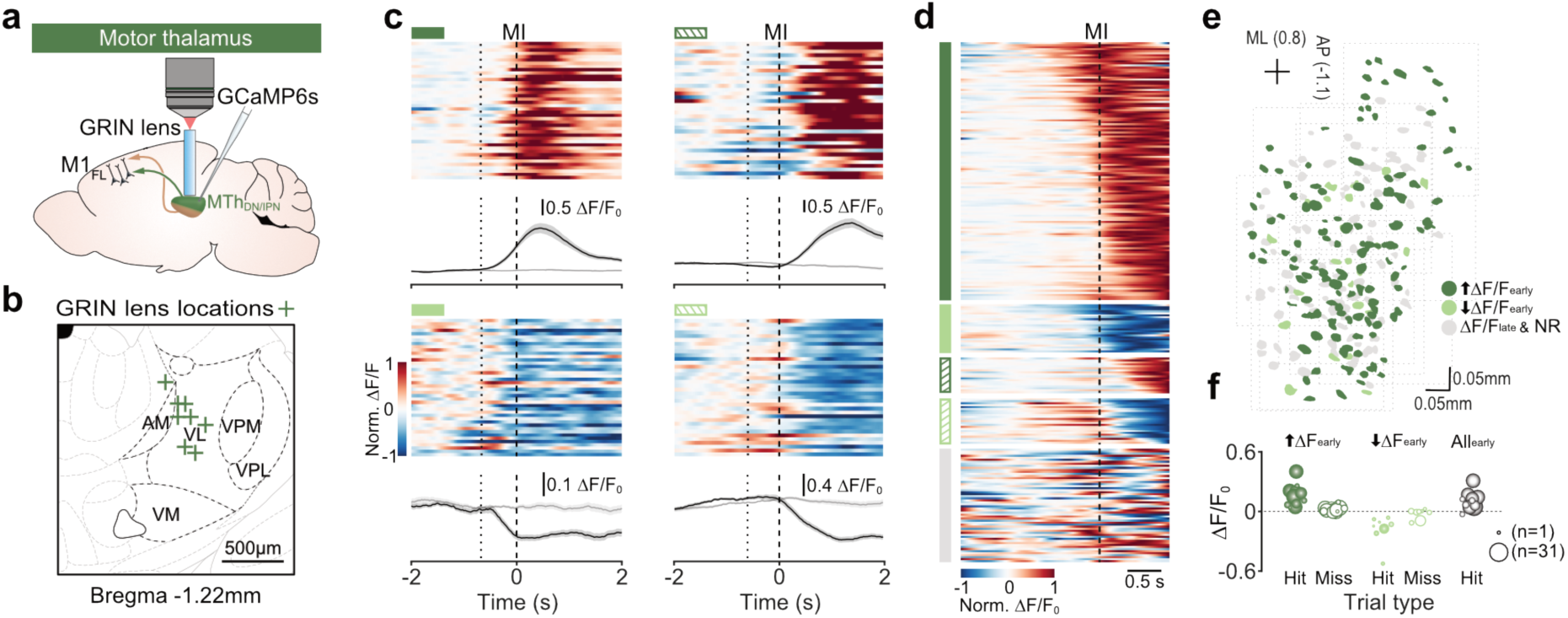
Enhanced activity dominates trial-to-trial MTh_DN/IPN_ population responses prior to movement initiation. **(a)** Gradient-index (GRIN) lens-mediated 2-photon population calcium imaging in MTh_DN/IPN_. M1_FL_, forelimb motor cortex; MTh_DN/IPN_ dentate / interpositus nucleus-recipient region of motor thalamus, **(b)** Anatomical locations of GRIN lens placements in MTh_DN/IPN_ (N = 8 mice). Motor thalamic nuclei: AM, anteromedial; VL, ventrolateral; VPM, ventral posteromedial nucleus; VPL, ventral posteromedial; VM, ventromedial, **(c)** Four example MThDN/IPN neurons: clockwise from top left, early onset enhanced’ (dark green), late onset enhanced’ (dark green hatching), ‘late onset suppressed’ (light green hatching), early onset suppressed’ (light green). *Top*, raster showing normalised ΔF/F_0_ across successive trials. Bottom, ΔF/F_0_ mean ± s.e.m. Black lines represent hit trials, grey lines represent miss trials. Dotted lines, median cue onset; dashed lines, movement initiation (MI), **(d)** Raster showing average ΔF/F_0_ across trials for individual neurons. Neurons are classified and grouped into early onset enhanced’ (dark green, n = 127/248 neurons); ‘early onset suppressed’ (light green, n = 24/248 neurons); late onset enhanced’ (dark green hatching, n = 18/248 neurons); late onset suppressed’ (light green hatching, n = 23/248 neurons); and ‘non-responsive’ (grey, n = 56/248 neurons). Neurons are ordered by ΔF/F_0_ onset (n = 11 fields of view. N = 8 mice), **(e)** Spatial distribution of early onset enhanced (dark green), early onset suppressed (light green) and late onset/non-responsive neurons (grey) in MTh_DN/IPN_. Dashed boxes represent different fields of view. ML, medial-lateral; AP anterior-posterior, **(f)** Average ΔF/F_0_ of early onset enhanced (dark green), early onset suppressed (light green) or all early onset (grey) neurons during hit (filled symbols) and miss trials (open symbols) separated by FOV. Circle size represents number of neurons per field of view [range 1 — 31].

We next tested which of our MTh_DN/IPN_ population response models best described the trial-to-trial activity in early onset enhanced neurons (see Figure 1b). Clustering trials by short, medium and long reaction times (RTs) and aligning averaged trial-to-trial population ΔF/F_0_ traces to movement initiation, MTh_DN/IPN_ responses displayed a consistent, sharp increase in ΔF/F_0_ ∼300 ms prior to movement initiation, irrespective of reaction time (short RT, mean onset = 308 ms, 95% CI [248, 377]; medium RT, mean onset = 299 ms, 95% CI [205, 392]; long RT, mean onset = 351 ms, 95% CI [251, 455], 117 neurons, 9 fields of view (FOV), N = 7 mice, for inclusion criteria see Methods) (Figures 3a-3d). Moreover, trial-to-trial consistency in MTh_DN/IPN_ output provided a reliable indication of when movement was likely to be initiated (Figures 3e and 3f). Thus, our results are consistent with a model whereby MTh_DN/IPN_ output provides a reliable time-locked motor signal immediately prior to movement initiation that is temporally uncoupled from the sensory cue (i.e. model (i) in Figure 1b).

**Figure 3.**
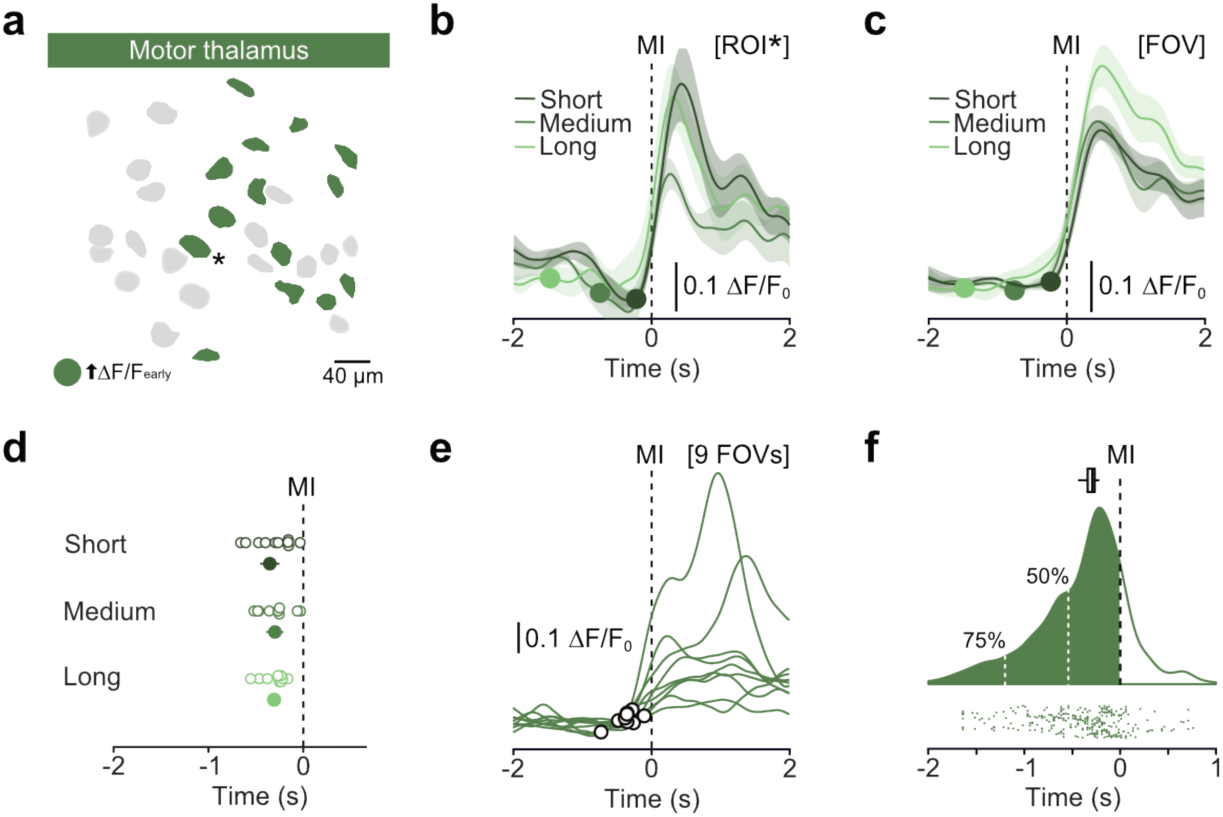
MTh_DN/IPN_ population responses provide a reliable trial-to-trial movement initiation signal. **(a)** Spatial distribution of early onset enhanced neurons in a representative field of view in MTh_DN/IPN_. **(b-c)** Average ΔF/F_0_ from (b) a single early onset enhanced neuron depicted in panel (a) by a *. or (c) all early onset enhanced neurons in the field of view shown in panel (a), aligned to movement initiation (MI) and split by short, medium and long reaction time trials. Colored circles depict median time of cue presentation. [ROI]. region of interest; [FOV], field of view. Mean ± s.e.m. **(d)** Motor thalamic population response trajectory onsets split by short, medium and long reaction times. Open circles represent individual fields of view, filled circles represent means ± 95% CI (n = 9 fields of view, N = 7 mice). Dashed line depicts movement initiation (MI), **(e)** Average ΔF/F_0_ trajectories from nine early onset enhanced neurons from different fields of view (FOVs) with response onsets indicated by black circles. Dashed line depicts movement initiation (MI), **(f)** Raincloud plot showing the distribution of bootstrapped trial-to-trial motor thalamic response onsets for all early onset enhanced neurons across nine fields of view. *Top*, box-and-whisker plot of the median onset bootstrapped estimate. *Middle*, average kernel density estimation of trial-to-trial motor thalamic response onsets with 50% and 75% AUC to movement initiation (MI) depicted by white vertical dashed lines. *Bottom*, raster of trial-to-trial early enhanced population onset times across all trials (n = 297 trials, n = 9 fields of view, N = 7 mice).

To gain a mechanistic insight into how feedforward input from MTh_DN/IPN_ shapes motor cortical membrane potential dynamics, we performed whole-cell patch-clamp recordings from identified L5B projection neurons in M1_FL_ (Figure 4a). When aligned to movement initiation, L5B neurons displayed a continuum of subthreshold membrane potential changes, biased towards depolarizing V_m_ (depolarizing, n = 15/23 neurons; hyperpolarizing, n = 4/23 neurons, non-responsive, n = 4/23, N = 23 mice), with the direction of the ΔV_m_ being consistent from trial-to-trial (Figures 4b-4f and Figure S5). Importantly, peak-scaled V_m_ traces – whether depolarizing or hyperpolarizing – displayed stereotyped trajectories where the ΔV_m_ onset closely matched the distribution of MTh_DN/IPN_ population response onsets (i.e. ∼300ms prior to movement initiation, Figures 4g-4i). To investigate whether L5B V_m_ dynamics are driven entirely by input from motor thalamus, we again exploited miss trials in which thalamic population responses are absent (see Figures 2c and 2f). V_m_ trajectories were on average smaller in amplitude and duration (mean miss:hit AUC ratio = 0.65, 95% CI [0.50, 0.80]), suggesting that convergence of thalamic and other long-range inputs is necessary for cued goal-directed movement initiation (Figure 4j and Figure S5). To drive movement, subthreshold V_m_ changes must transform into behaviorally-relevant spiking. Accordingly, we found a strong correlation between changes in V_m_ and firing rate across layer 5B projection neurons, including pyramidal tract (PT-type) neurons that have direct access to brainstem and spinal cord circuits controlling voluntary movement (Figures 4k-4m and Figure S5). Next, we focally injected a small bolus of muscimol centered on the ventrolateral nucleus while recording from identified L5B projection neurons (Figure 4n) to explore a causal link between thalamic input, L5B V_m_ trajectories and movement initiation. Muscimol inactivation reduced the amplitude and duration of L5B V_m_ responses (mean muscimol:control AUC ratio = 0.35, 95% CI [0.26, 0.44], n = 3 neurons from N = 3 mice), mirroring V_m_ trajectories during miss trials where MTh_DN/IPN_ input is absent (compare Figure 4j and Figure 4p; and Figure 4o and Figure S5h), and reduced task success (mean = 0.37 normalized task success, 95% CI [0.16, 0.59], n = 3 neurons, N = 3 mice) (Figure 4q). Thus, MTh_DN/IPN_ input to M1_FL_ drives activity dynamics necessary for goal-directed movement initiation.

**Figure 4.**
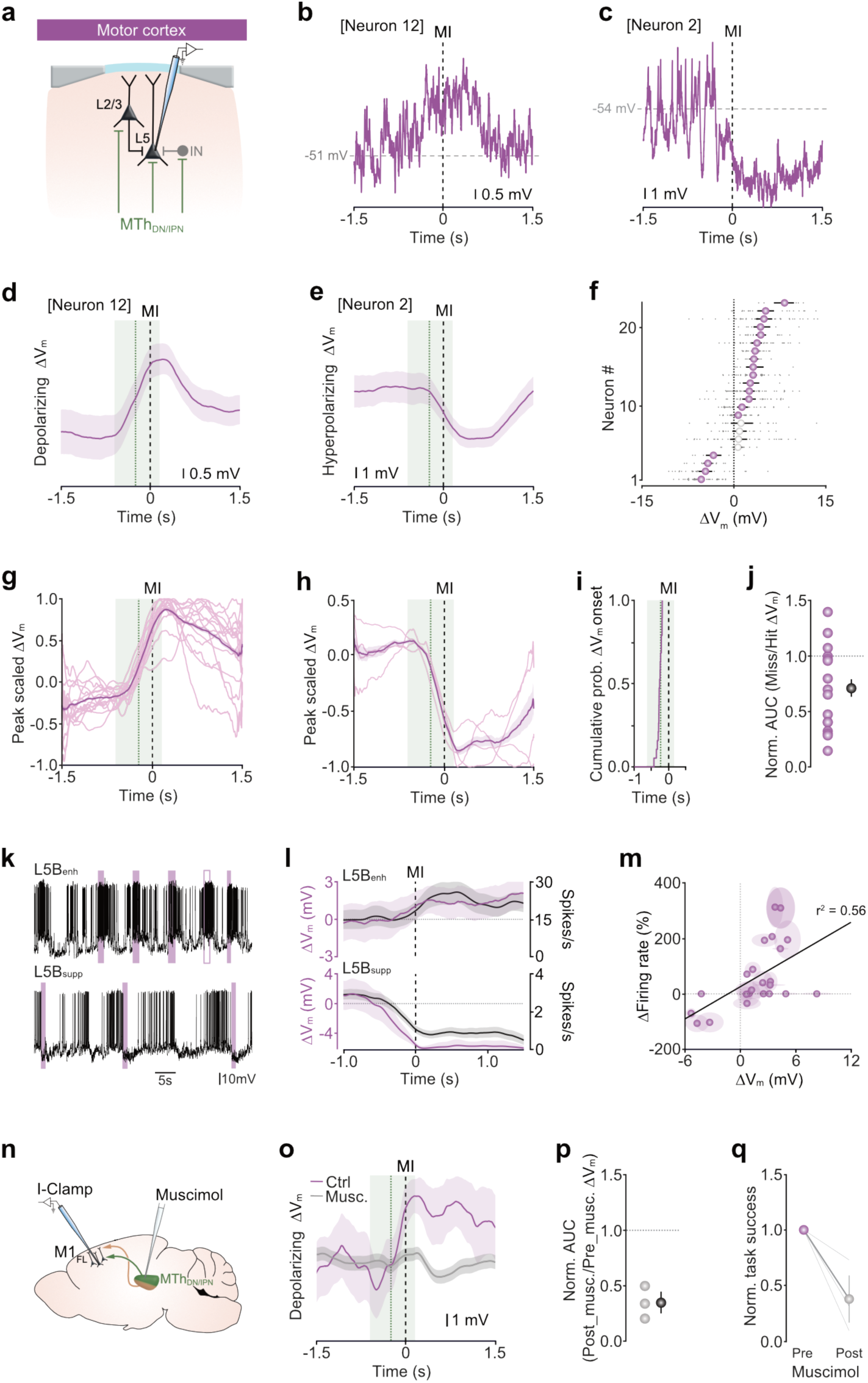
Feedforward MTh_DN/IPN_ input is necessary for bidirectional M1_FL_ L5B output modulation and cued goal-directed movement initiation. **(a)** Patch-clamp recording from L5B projection neurons in M1_FL_ IN, interneuron; MTh_DN/IPN_, dentate / interpositus nucleus-recipient region of motor thalamus, **(b-c)** Representative single trial subthreshold membrane potential (V_m_) trajectories from two L5B projection neurons showing either a depolarization (b) or hyperpolarization (c) prior to movement initiation (MI). Spikes have been clipped to improve visualisation of the subthreshold V_m_. **(d-e)** Average changes in subthreshold V_m_ ± 95% CI in the two L5B projection neurons depicted in (b) and (c). In these and subsequent figure panels, the green dashed line depicts mean MTh_DN/IPN_ activity onset ± 95% CI as shown in Fig. 3d and the black dashed line represents movement initiation (MI), **(f)** Average L5B projection neuron perimovement ΔV_m_ ± 95% CI. Grey dots represent individual trials, purple symbols represent significant ΔV_m_ changes, white symbols represent non-significant changes, defined by comparing 95% bootstrapped confidence intervals (n = 23 neurons, N = 23 mice), **(g-h)** Peak scaled mean subthreshold V_m_ from individual neurons overlaid and split by direction of change (g, depolarizing, n = 15/23 neurons; h, hyperpolarizing, n = 4/23 neurons). Thick purple line represents population means ± 95% CI. **(i)** Cumulative probability of ΔV_m_ onsets across all significantly modulated L5B projection neurons (n = 19/23 neurons). MI, movement initiation, **(j)** Ratio of normalised area under the curve for V_m_ trajectories during miss versus hit trials. Purple symbols represent data from individual neurons, black symbols represent population mean ± 95% CI. **(k)** Representative V_m_ traces from a L5B depolarizing (top) and L5B hyperpolarizing (bottom) neuron across multiple trials. Filled purple bars depict hit trials, open purple bars depict miss trials. For clarity, internally generated movements (IGMs) are not shown. **(I)** Average subthreshold ΔV_m_ (purple) and firing rate (FR, black) trajectories for the L5B enhanced and L5B suppressed neurons shown in (k) aligned to movement initiation (MI). Thick lines represent the mean ± 95% CI. **(m)** Correlation between movement-related subthreshold ΔV_m_ and firing rate changes. Colored symbols represent mean ± 95% CI from individual neurons, black line is a linear fit to the data (Pearson’s *r*). **(n)** Patch-clamp recording from M1_FL_ L5B projection neurons during muscimol inactivation targeted to MTh_DN/PN_. I-Clamp, current clamp; M1_FL_, forelimb motor cortex, **(o)** Average subthreshold ΔV_m_ ± 95% CI from a L5B projection neuron before (Ctrl, purple) and after muscimol inactivation (Muse., black) targeted to MTh_DN/PN_. **(p)** Ratio of normalised area under the curve for V_m_ trajectories post (Post_musc.) versus pre (Pre_musc.) muscimol injection. Grey symbols represent data from individual neurons, black symbol represents population mean ± 95% CI. **(q)** Normalised task success before (purple symbol) and after (grey symbol) muscimol injection targeted to MTh_DN/IPN_. Colored symbols represent mean ± 95% Cl. grey lines indicate data from individual neurons.

If prior to movement, mice remain in a prepared state awaiting an ‘initiation’ signal, direct stimulation of the MTh_DN/IPN_ thalamocortical pathway in the absence of the cue could provide an input sufficient to evoke learned movements. We tested this prediction using a dual optogenetic stimulation strategy employing either direct stimulation of MTh_DN/IPN_ neurons or thalamocortical axon terminals in M1_FL_ during the baseline period prior to cue presentation (Figure 5a). By targeting small volumes of AAV2/1-CAG-ChR2 virus to the dorsal-posterior motor thalamus, we restricted opsin expression almost exclusively to neurons in MTh_DN/IPN_ (Figure S6). Direct stimulation of MTh_DN/IPN_ or axon terminals in M1_FL_ evoked forelimb movements in 9/10 mice, with full lever pushes occurring in ∼26% of trials (MTh_DN/IPN_ stimulation, mean = 0.27 proportion of trials with full lever push, 95% CI [0.23, 0.30], N = 9/10 mice; axon terminal stimulation, mean = 0.25 proportion of trials with full lever push, 95% CI [0.13, 0.38], N = 6/6 mice, Video S4). The duration and reaction times of photostimulated push movements were comparable to cue-evoked trials (Figures 5b-5d). In a small proportion of trials, light stimulation evoked partial lever pushes that did not reach the reward zone (MTh_DN/IPN_ stimulation, mean = 0.15 proportion of trials, 95% CI [0.09, 0.20], N = 9 mice; axon terminal stimulation, mean = 0.06 proportion of trials, 95% CI [0, 0.14], N = 6 mice) (Figure 5b). Stimulating either pathway in the absence of ChR2 expression did not evoke any detectable forelimb movements (N = 2) (Figure S6). To compare the cellular effects of cueversus ChR2-evoked MTh_DN/IPN_ input in M1_FL_, we performed whole-cell recordings from identified M1_FL_ L5B projection neurons during interleaved cue and MTh_DN/IPN_ photostimulation trials (Figure 5e). Remarkably, during photostimulation trials, full push movements were associated with depolarizing or hyperpolarizing V_m_ changes that matched cue-evoked V_m_ changes in the same neuron (Figures 5f-5h). Thus, selective recruitment of the MTh_DN/IPN_ – M1_FL_ thalamocortical pathway can recapitulate M1_FL_ L5B neural activity dynamics required for goal-directed movement initiation.

**Figure 5.**
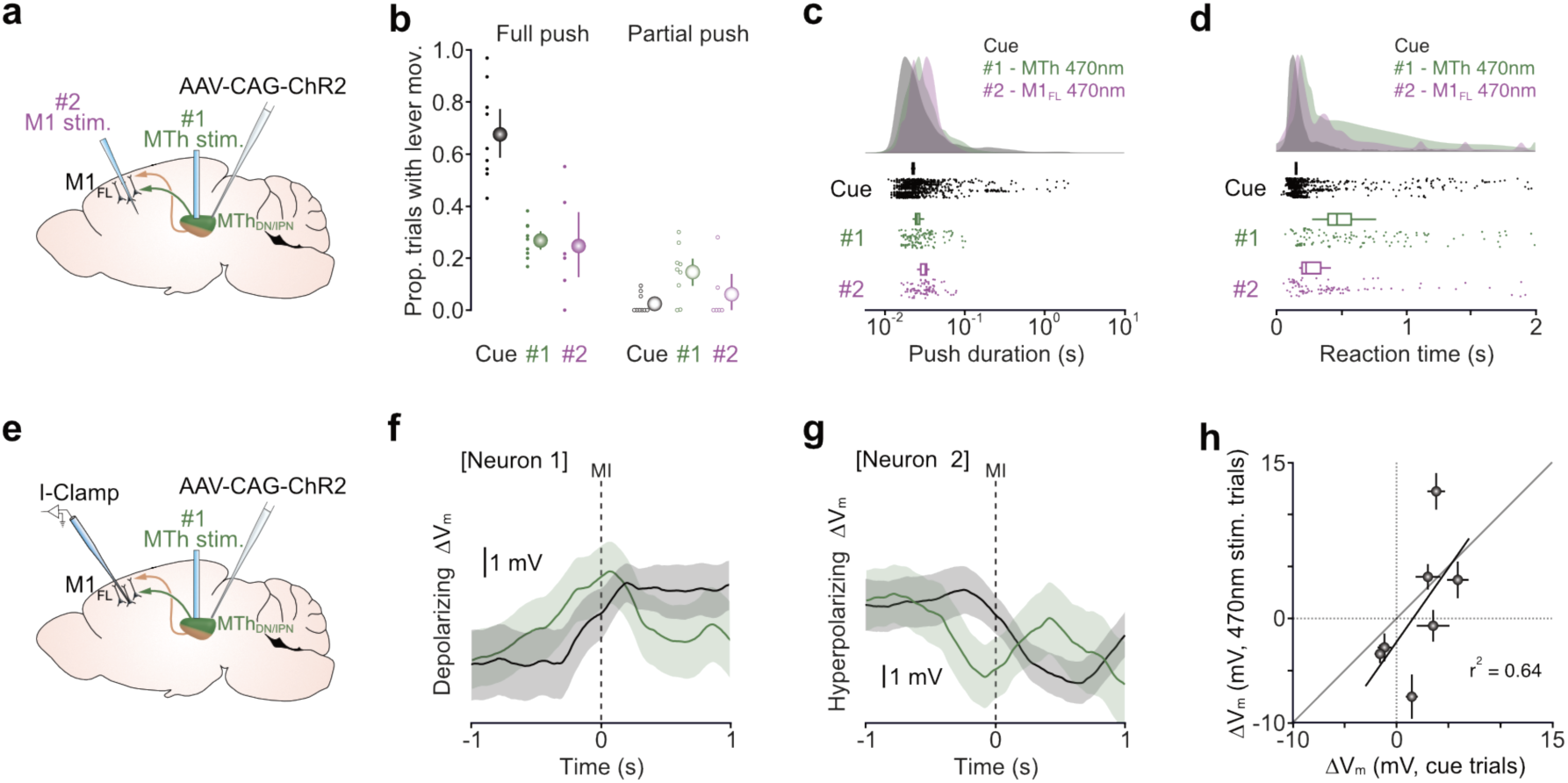
Optogenetic stimulation of MTh_DN/IPN_ axon terminals recapitulates cued goal-directed movement. **(a)** Dual MTh optogenetic stimulation strategy: Channelrhodopsin 2 (ChR2) expression was targeted to neurons in MTh_DN/IPN_ and stimulated via an optic fiber placed directly above (#1) or via a tapered optic fiber implanted directly into forelimb motor cortex (M1_FL_) (#2). **(b)** Comparison of cue-evoked and ChR2-evoked task engagement represented as the proportion of trials with either full (*left*) or partial (*right*) lever push movements. Black, cue-evoked: green, direct MTh_DN/IPN_ stimulation: purple, stimulation of MTh_DN/IPN_ axon terminals in M1_FL_. Colored dots represent data from individual mice, colored symbols represent mean ± 95% CI. For Cue. #1 and #2. N = 9. 9 and 6 mice, respectively, **(c-d)** Raincloud plots showing the distributions of push durations (c) and reaction times (d) of cue-evoked (black) and ChR2-evoked (#1. green and #2. purple) hit trials. Box-and-whisker plots of bootstrapped estimates of median statistics, **(e)** Patch-clamp recording from M1_FL_ L5B projection neurons during ChR2-mediated stimulation of MTh_DN/IPN_ neurons. I-Clamp, current clamp; M1_FL_, forelimb motor cortex, **(f-g)** Average changes in subthreshold V_m_ ± 95% CI in two L5B projection neurons showing either a pre-movement depolarization (f) or hyperpolarization (g) in response to an auditory cue (black) or ChR2-mediated stimulation of MTh_DN/IPN_ neurons (green). Dashed line represents movement initiation, **(h)** Correlation between peri-movement cue-evoked and ChR2-evoked subthreshold Δ V_m_ across L5B projection neurons (n = 7 neurons, N = 6 mice). Filled symbols represent mean ± 95% CI, black line is a linear fit to the data (Pearson’s *r*).

If behavioral context provides information necessary for learned movement initiation, ChR2-evoked forelimb lever push movements should be abolished during direct MTh_DN/IPN_ stimulation in an altered behavioral context. To test this prediction, we placed trained mice on a flat baseplate in the absence of any support / movable levers, reward spout or water rewards, and compared cue-evoked forelimb movements in both the learned and altered behavioral contexts. Habituation in the altered behavioral context was performed within training session to ensure that the cued lever push behavior was not extinguished. As expected, trained mice generated cue-evoked forelimb lever push trajectories in 63% of trials in the learned behavioral context (LBC) but in the altered behavioral context (ABC) cue-evoked push-like movements were absent (LBC, mean = 0.63 proportion of trials, 95% CI [0.55, 0.71]; ABC, mean = 0.02 proportion of trials, 95% CI [0.00, 0.04], N = 3 mice), confirming that mice acknowledged the difference between the two behavioral environments (Figures 6a-6e, Video S5). We then replicated the experiment by replacing the auditory cue with direct photostimulation of MTh_DN/IPN_ during the baseline period (Figure 6f). If direct MTh_DN/IPN_ stimulation alone drives specific muscle synergies, then ChR2-evoked forelimb movement trajectories will be unaffected by a change in context. However, we found that in the learned behavioral context, direct MTh_DN/IPN_ stimulation evoked forelimb movements in 67% of trials with 45% of trials containing successful lever push trajectories (mean = 0.67 proportion of trials with movement, 95% CI [0.52, 0.82]; mean = 0.45 proportion of trials with push action, 95% CI [0.40, 0.49], N = 3 mice) (Figures 6g and 6i, Video S6). While in the altered behavioral context, direct MTh_DN/IPN_ stimulation evoked forelimb movements in 64% of trials but only 4% contained push-like movements (mean = 0.64 proportion of trials with movement, 95% CI [0.59, 0.69]; mean = 0.04 proportion of trials with push action, 95% CI [0.02, 0.07], N = 3 mice) (Figures 6h and 6j, Video S6). Taken together, these results demonstrate that the MTh_DN/IPN_ thalamocortical pathway conveys a robust motor timing signal necessary for initiating behavioral context-specific movement initiation.

**Figure 6.**
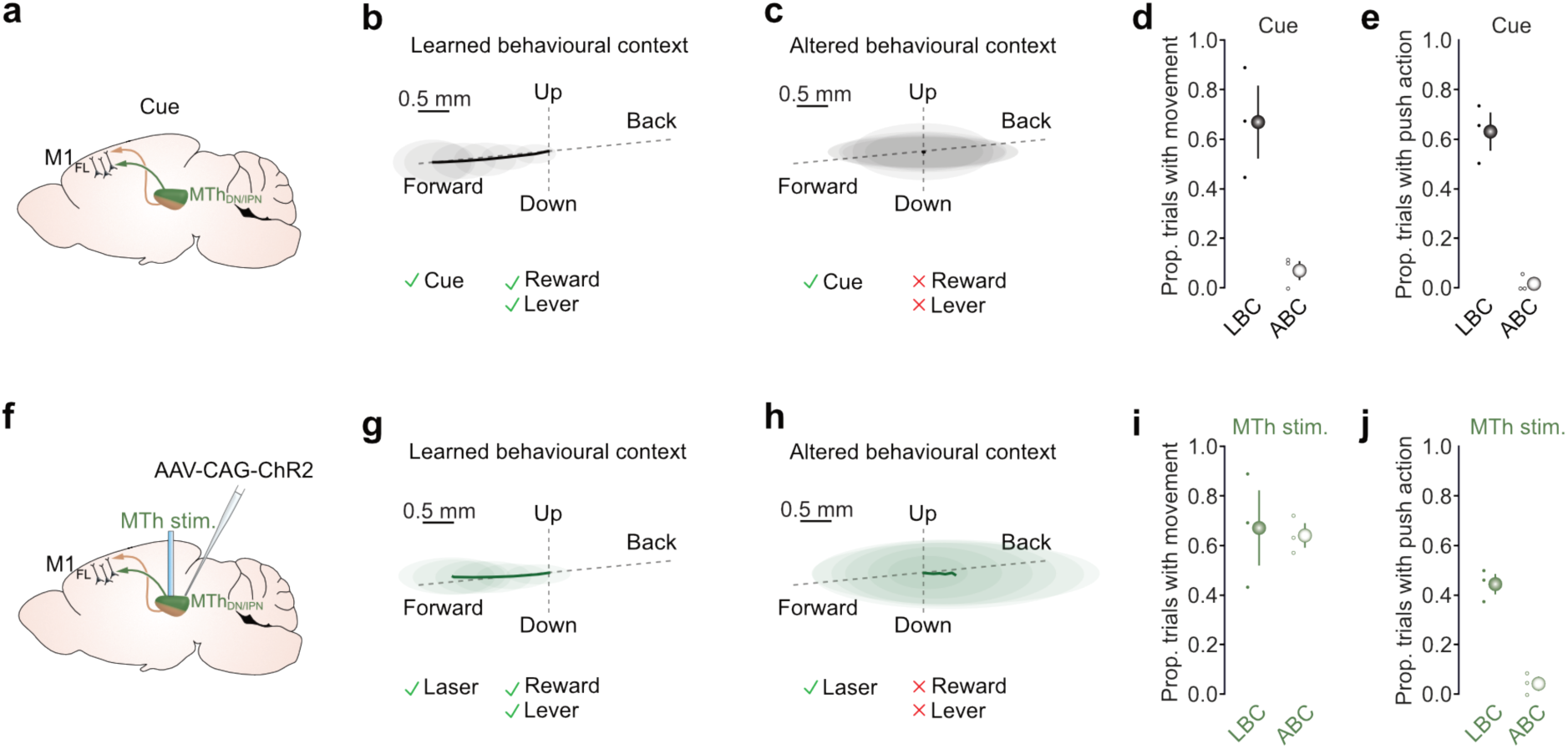
MTh_DN/IPN_ stimulation evokes behavioral context-specific movement initiation. **(a)** Mouse sagittal brain schematic depicting cue-evoked feedforward input from MTh_DN/IPN_,to M1_FL_. **(b-c)** Average cue-evoked kinematic forepaw trajectories from an example mouse in a learned behavioral context (i.e. auditory cue, reward spout & movable lever (b) or altered behavioral context (i.e. auditory cue, no reward spout & no movable lever (c). Thick black line depicts average trajectory overlaid with the 95% CI of frame-by-frame paw position variance, **(d-e)** Proportion of trials with cue-evoked forelimb movement (d) or forelimb push actions (e) in a learned behavioral context (LBC) versus altered behavioral context (ABC). Colored dots represent data from individual mice, colored symbols represent mean ± 95% CI (N = 3 mice), **(f)** MTh_DN/IPN_ optogenetic stimulation strategy: ChR2 expression was targeted to neurons in MTh_DN/IPN_ and stimulated via an optic fiber placed directly above, **(g-h)** Average ChR2-evoked kinematic forepaw trajectories from an example mouse in a learned behavioral context (g) or altered behavioral context (h). Thick green lines depict average trajectory overlaid with the 95% CI of frame-by-frame paw position variance, **(i-j)** Proportion of trials with ChR2-evoked forelimb movement (i) or forelimb push actions (j) in a learned behavioral context (LBC) versus altered behavioral context (ABC). Colored dots represent data from individual mice, colored symbols represent mean ± 95% CI.

## Discussion

Here, we investigated the contribution of the cerebellar-recipient motor thalamocortical pathway to movement initiation, showing that a robust and reproducible feedforward motor timing signal propagating from MTh_DN/IPN_ to M1_FL_ is essential for goal-directed movement initiation. Specifically, we show that trial-to-trial MTh_DN/IPN_ population responses are dominated by a time-locked increase in activity immediately prior to movement initiation that is temporally uncoupled from cue presentation, providing a fixed latency feedforward motor timing signal to M1_FL_. MTh_DN/IPN_ thalamocortical input is a prerequisite for generating M1_FL_ layer 5B activity dynamics necessary for movement initiation and blocking MTh_DN/IPN_ output suppresses task engagement. Finally, direct stimulation of MTh_DN/IPN_, or their axon terminals in M1_FL_, in the absence of the cue recapitulated motor cortical activity dynamics and forelimb behavior in the learned behavioral context, but generated semi-random movements in an altered behavioral context where the lever and reward were absent. Together, these data suggest that dentate and interpositus nucleus-recipient motor thalamocortical pathway plays a pivotal role in directly gating movement initiation, thus confirming and extending existing theories of the role of the cerebellar-thalamocortical pathway in initiating goal-directed movement.

By employing population calcium imaging of MTh_DN/IPN_ activity we demonstrate that trial-to-trial output from the dentate and interpositus nucleus-recipient regions of motor thalamus do not reflect sensorimotor transformations from cue to movement initiation, instead we suggest that MTh_DN/IPN_ output reflects a pure feedforward motor timing signal that indicates the immediate intention to move. In the absence of this input, i.e. local inactivation of MTh_DN/IPN_, the command to move is blocked resulting in a suppression of goal-directed movement initiation. If the cerebellar-recipient motor thalamocortical pathway conveys a pure motor timing signal, where in the brain is the delay between cue and MTh_DN/IPN_ activity onset generated? A signal that directly gates movement likely overlaps with preparatory activity in frontal motor regions (Churchland et al., 2006b; Li et al., 2015; Requin et al., 1990) irrespective of how movements are initiated (Lara et al., 2018). Preparatory activity both in frontal motor regions and deep cerebellar nuclei are driven by a cortico-cerebellar loop through the motor thalamus, where persistent neural dynamics across brain regions facilitates movement choice and execution. Thus, sensory-driven persistent activity in frontal motor-associated regions could provide the initial ‘decision to move’, which propagates through cortico-cerebellar loops to form a discrete motor timing signal at the level of motor thalamus (Chabrol et al., 2019; Churchland et al., 2006a; Gao et al., 2018; Guo et al., 2014). Although our data do not shed light on the origin of the signal, we demonstrate that MTh_DN/IPN_ is an essential node in the cerebello-cortical loop through which motor timing signals propagate to initiate goal-directed movement. Further studies will be required to determine the contribution of cortico-ponto-cerebellar and cerebellar-thalamocortical loops to sensorimotor transformations across a range of goal-directed motor tasks, and whether information pertaining to movement preparation and execution propagate through the same or parallel subdivisions of the motor thalamus (Chabrol et al., 2019; Gao et al., 2018; Kuramoto et al., 2009; Miller and Brooks, 1982).

The behavioral context-dependence of photoactivated movements suggests that MTh_DN/IPN_ likely conveys a movement-invariant motor initiation signal that converges, at the level of motor cortex, with other long-range inputs necessary for selecting movement type. Consistent with this notion, in the absence of feedforward MTh_DN/IPN_ input (i.e. during miss trials or thalamic inactivation), M1 layer 5 projection neurons displayed reduced task-related activity that did not initiate movement. The origin of the convergent long-range input(s) remains unknown, but likely candidates are cortico-cortical interactions between orbitofrontal cortex or frontal motor areas and M1 (Hooks et al., 2013; Reep et al., 1990), thought to accumulate task-relevant information required for motor planning and decision-making (Gao et al., 2018; Li et al., 2015), or basal ganglia-thalamocortical interactions that determine the type, timing and invigoration of upcoming movements (Dudman and Krakauer, 2016; Inase et al., 1996; Klaus et al., 2019; Thura and Cisek, 2017; Williams and Herberg, 1987).

Directly activating the MTh_DN/IPN_ thalamocortical pathway in the altered behavioral context consistently generated semi-random forelimb movements (Tanaka et al., 2018), whereas full recapitulation of the learned behavior could only be achieved in the learned behavioral context. If convergent input from MTh_DN/IPN_ and other task-related brain areas is necessary for learned movement initiation, why can photostimulation of the thalamocortical pathway result in learned movement initiation in the absence of an external sensory cue? Previous studies have shown that behavioral context is an important determinant of neural trajectories during goal-directed motor tasks (Russo et al., 2018; Suresh, 2019), where cortical dynamics evolve in a pre-determined manner depending on their initial state (Churchland et al., 2010; Kaufman et al., 2014; Sauerbrei, 2018). In the learned behavioral context, M1 population dynamics likely remain in a quasi-prepared state, awaiting external input to initiate learned forelimb movement. Thus, in some trials (up to 40%) direct MTh_DN/IPN_ stimulation is likely sufficient to drive M1 neural trajectories towards a state required for learned movement initiation. Conversely, in mice habituated to an unrewarded, altered behavioral context, M1 population dynamics driven by direct MTh_DN/IPN_ stimulation likely evolve from a different initial state resulting in neural trajectories that generate arbitrary, but not goal-directed, forelimb movements (Graziano et al., 2002; Rispal-Padel et al., 1982; Tanaka et al., 2018). A central question for future investigation is how thalamic input and behavioral context contribute to motor cortical population dynamics across learning and different motor behaviors.

In summary, our findings extend our understanding of how specific subdivisions of the mammalian motor thalamus contribute to motor timing (Dormont et al., 1982; Kurata, 2005; Strick, 1976), suggesting that the cerebellar-thalamocortical pathway plays a critical role in the initiation of goal-directed movement.

## Acknowledgements

We are grateful to Tiago Branco, Benjamin Grewe, Jan Gründemann, Matthew Nolan, Gülsen Sürmeli, Brett Mensh and members of the Nolan and Duguid labs for experimental discussions and for comments on the manuscript. Nick Steinmetz for the suggested design of the author contribution matrix. Pseudotyped SADΔG-mCherry(EnvA) rabies virus was a generous gift from Edward Callaway (Salk Institute for Biological Studies) to A.H. AAV-GCaMP6s was a gift from Douglas Kim & GENIE Project (Addgene 100844-AAV1). pACAGW-ChR2-Venus-AAV was a gift from Karel Svoboda (Addgene plasmid #20071) and packaged by Christina McClure and Innes Jarmson (Nolan Lab, University of Edinburgh). Confocal microscopy was performed in the IMPACT Imaging Facility at the University of Edinburgh. We thank Marie Zechner for assistance with graphic design. Research was supported by grants from the Biotechnology and Biological Sciences Research Council (BB/R018537/1), DFG fellowship program (SCHI1267/2-1 to J.S. and AM 443/1-1 to J.A.), the Shirley Foundation, Howard Hughes Medical Institute (A.H. and C-C.H.) and a Wellcome Senior Research Fellowship (110131/Z/15/Z) to I.D.

## Author contributions

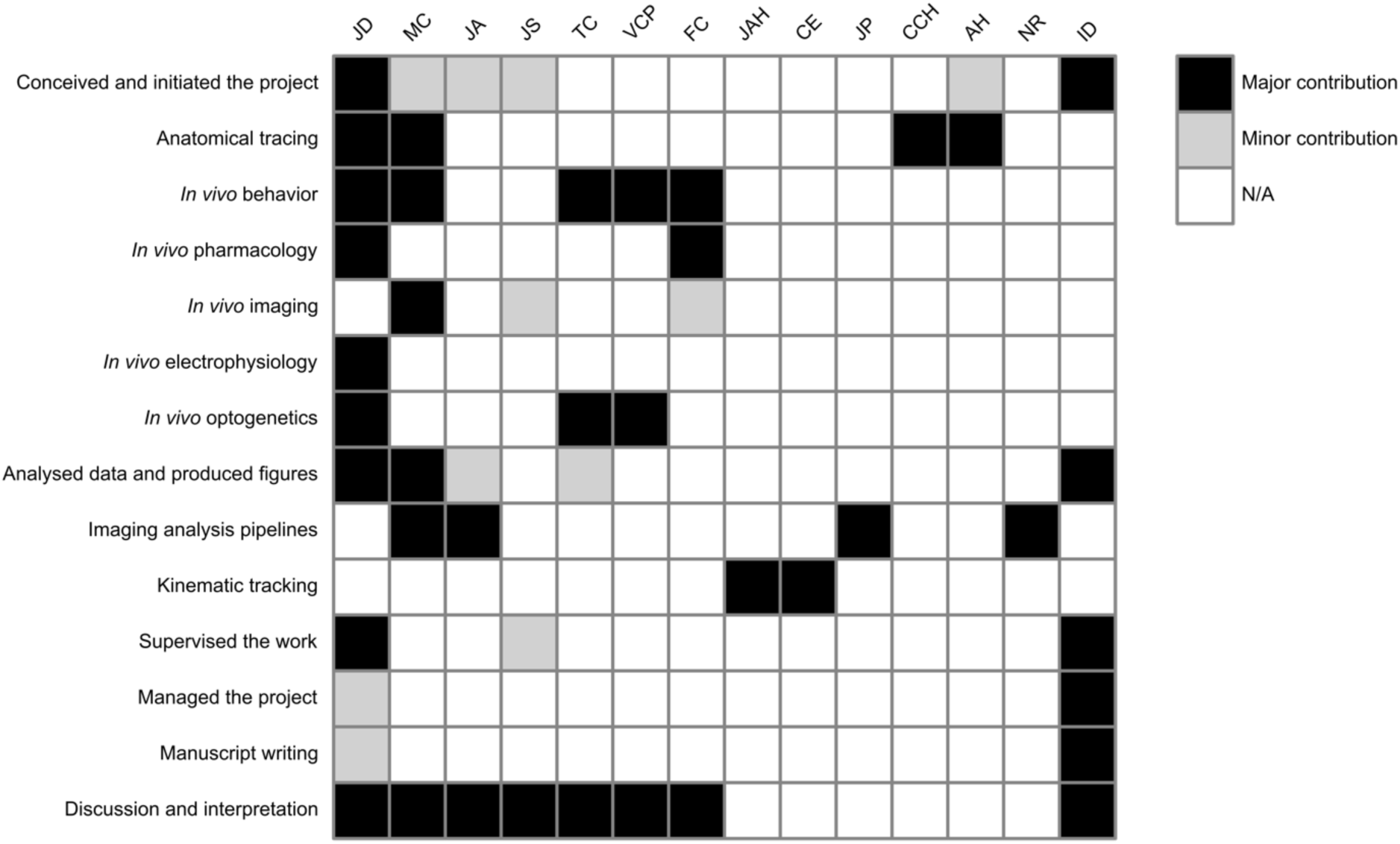

## Methods

### Animal husbandry and general surgery

Male adult C57BL/6J wild-type and Rbp4-Cre (MMRRC, stock 031125-UCD) mice (5-12 weeks old, 20-30g, one to six animals per cage) were maintained on a reversed 12:12 hour light:dark cycle and provided *ad libitum* access to food and water. All experiments and procedures were approved by the University of Edinburgh local ethical review committee and performed under license from the UK Home Office in accordance with the Animal (Scientific Procedures) Act 1986. Surgical procedures were performed under ∼1.5% isoflurane anesthesia and each animal received fluid replacement therapy (0.5ml sterile Ringer’s solution) to maintain fluid balance and buprenorphine (0.5 mg/kg) for post-operative pain relief. A small lightweight headplate (0.75g) was implanted on the surface of the skull using cyanoacrylate super glue and dental cement (Lang Dental, USA) and mice were left for 24-48 hours to recover. Craniotomies were performed in a stereotactic frame (Kopf, USA) using a hand-held dentist drill with 0.5 mm burr (whole-cell patch-clamp recording ø300 µm; viral/tracer/pharmacological compound injection ø500-1000 µm), viral vectors and tracing compounds were delivered via pulled glass pipettes (5μl, Drummond, 10-20 nl/min) using an automated injection system (Model Picospritzer iii, Intracell).

### Monosynaptic retrograde rabies tracing

For monosynaptic retrograde rabies tracing, conditional expression of TVA receptor was achieved by injecting 60nl of AAV2/1-CAG-FLEX-mTagBFP2-2A-TVA (9.0×10^12^ genome copies per ml (GC/ml)) into contralateral M1_FL_ (AP: 0.6, ML: 1.6, DV: -0.7 mm) of three Rbp4-Cre mice. For anterograde labelling of deep cerebellar nuclei projections to MTh_DN/IPN_, AAV2/1-CAG-EGFP (1.1×10^13^ GC/ml) was vertically injected into contralateral dentate (AP: -6.2, ML: 2.25, DV: -2.5 & -2.0 mm) and interpositus deep cerebellar nuclei (AP: -6.2, ML: 1.25, DV: 2.5 & -2.0 mm), with 60 nl injected at each depth. Pseudotyped SADΔG-mCherry(EnvA) rabies virus (produced as previously described (Wickersham et al., 2007; Wickersham et al., 2010) was injected into M1_FL_ (same coordinates as stated previously) three weeks after the initial injections. Mice were perfused seven days post-rabies virus injection. Sections (60 μm) were cut, mounted and imaged at 20x using a Nanozoomer Slide Scanner (Hamamatsu). For all anatomical quantification, raw data images were manually referenced to the Paxinos & Franklin Mouse Brain Atlas (Paxinos & Franklin, 2008). The distribution of fluorescence was manually outlined and independently verified.

### Conventional retrograde tracing

For retrograde tracing of M1_FL_-projecting motor thalamic neurons, a single (∼ø1 mm) craniotomy was performed above contralateral M1_FL_ (AP: 0.6, ML: 1.6, DV: -0.7 mm), and 150 nl of red (590 nm) retrobeads (Lumafluor Inc.) was injected at four points equidistant from the center of the craniotomy. After recovery, mice were returned to the home cage for ∼7 days before being anaesthetized with euthatal (0.10–0.15 ml) and transcardially perfused with 30 ml of ice-cold PBS followed by 30 ml of 4% paraformaldehyde (PFA) in PBS solution. Brains were post-fixed in PFA overnight at 4°C and transferred to 10% sucrose solution for storage. Coronal sections (60 μm) were cut with a vibratome (Leica VT1000S), mounted using Vectashield mounting medium (H-1000, Vector Laboratories), and imaged using a laser scanning confocal microscope (Leica TCS-NT). To assess the density of M1_FL_-projecting neurons originating in the ventrolateral motor thalamus, 200nl of CTB-Alexa647 (Invitrogen) was injected into M1_FL_ (AP: 0.6, ML: 1.6, DV: -0.7 mm). After 7 days post injection, mice were perfused (as described above) and 100µm sections were counter-stained with Nissl blue before being imaged with a Leica LSM800 confocal microscope. Cells were counted in a representative 300 × 300 µm region of the ventrolateral motor thalamus and counts were independently verified.

### Behavioral training

After recovery from head plate surgery, mice were handled extensively before being head restrained and habituated to a custom lever push behavioral setup. Mice were trained to perform horizontal lever push movements (4 mm) in response to a 6kHz auditory cue in order to obtain a 4-8 µl water reward. To increase task engagement mice were placed on a water control paradigm (1 ml/day) and weighed daily to ensure body weight remained above 85% of baseline. Mice were trained once per day for 30 mins, with a quasi-random inter-trial-interval of 4-6s followed by presentation of an auditory cue. Mice responded within a 10 s window early in training, reduced to a 2 s window prior to recording, and were deemed ‘expert’ after achieving >90 rewards per session on two consecutive days. Lever movements during the ITI would result in a ‘lever reset’ and commencement of a subsequent ITI.

### *In vivo* pharmacology

To assess the behavioral effects of M1_FL_ / MTh_DN/IPN_ inactivation, the contralateral forelimb was shaved under general anesthesia and the wrist, elbow and shoulder joints were marked with black ink. Mice were allowed to recover for at least 60 mins before being head-restrained in the behavioral apparatus. After 5 min of baseline task execution, the lever was locked and a small volume of the GABA_A_ receptor agonist muscimol (dissolved in external solution containing 150 mM NaCl, 2.5 mM KCl, 10 mM HEPES, 1.5 mM CaCl_2_ and 1 mM MgCl_2_) or saline was injected into the target area (M1_FL_: 200 nl of 2 mM muscimol at each of 5 sites centered on AP: 0.6, ML: 1.6, DV: -0.7 mm; MTh_DN/IPN_: 200 nl of 1 mM muscimol, AP: -1.1, ML: 1.0, DV: -3.4 mm). Mice were randomly assigned to drug or control groups, and experiments performed blinded. To confirm anatomical location of each drug injection, 1% w/v of red (590 nm) retrobeads (Lumaflor Inc.) was included in the drug/saline solution. Behavioral metrics were analyzed in 5-minute epochs using a two-way repeated measures ANOVA to determine statistical significance with Bonferroni-Holm correction for multiple comparisons.

### GRIN lens imaging

To perform population calcium imaging in motor thalamus we injected 200 nl of AAV1-Syn-GCaMP6s (2.9×10^13^ GC/ml, Addgene 100844-AAV1) into contralateral MTh_DN/IPN_ (AP: -1.1, ML: 1.0, DV: -3.4 mm) before implanting a lightweight headplate as described above. After 7-10 days, a gradient-index (GRIN) lens (Grintech NEM-060-15-15-520-S-1.0p; 600 μm diameter, 4.83 mm length, 0.5 numerical aperture) was implanted as described previously (Xu et al., 2016). In brief, a sterile needle (1.1 mm OD) surrounding the GRIN lens was lowered to a depth of 3.2 mm and subsequently retracted leaving the lens at the desired depth. The lens was then secured in place with UV curing glue (Norland Products, USA) and dental cement (Lang Dental, USA). Fields of view were checked every 14 days for clarity and GCaMP6s expression. After 4-8 weeks mice began water restriction and behavioral training. Two-photon calcium imaging was performed in expert mice during task engagement with a 320 x 320 µm field of view (600 x 600 pixels) at 40 Hz frame rate, using a Ti:Sapphire pulsed laser (Chameleon Vision-S, Coherent, CA, USA; < 70 fs pulse width, 80 MHz repetition rate) tuned to 920 nm wavelength with a 40x objective lens. For confirmation of GRIN lens location and viral expression, mice were perfused as described above and sections (100 μm) were cut with a vibratome, counterstained with Nissl blue, and imaged using a slide scanner (Zeiss Axioscan). GRIN lens location was determined using the Paxinos & Franklin Mouse Brain atlas (Paxinos & Franklin, 2008), and anatomical confirmation within MTh_DN/IPN_ was used to determine data inclusion. Motion artefacts in the raw fluorescence videos were corrected using NoRMCorre (Pnevmatikakis et al., 2017). In brief, NoRMCorre performs non-rigid motion correction by splitting each FOV into overlapping patches, estimating the xy translation for each patch, and upsampling the patches to create a smooth motion field, correcting for non-uniform motion artefacts caused by raster scanning or brain movement. Regions of interest (ROIs, polygonal areas) were manually drawn in Fiji (Schindelin et al., 2012) and fluorescence signals were decontaminated and extracted using nmf_sklearn to remove fluorescence originating from neuropil and neighboring cells (Keemink et al., 2018). Normalized fluorescence was calculated as ΔF/F_0_, where F_0_ was calculated as the 5th percentile of the 1Hz low-pass filtered raw fluorescence signal and ΔF = F-F_0_. To define early responsive neurons, average ΔF/F_0_ signals during baseline and peri-movement epochs were compared (baseline epoch = 500 ms pre-cue; movement epoch = -250 to +500 ms peri-movement) using a Wilcoxon rank sum test with a significance threshold of *P*<0.01. The direction of the response was defined as suppressed or enhanced by comparing the median value of the ΔF/F_0_ signal during both epochs. Late responsive suppressed/enhanced neurons were identified by comparing the 500 ms pre-cue baseline epoch with a 500 ms pre-reward epoch using a Wilcoxon rank sum test with a significance threshold of *P*<0.01. For presentation, movement-aligned ΔF/F_0_ signals were smoothed with the loess method using a 40-frame sliding window and baseline corrected to the mean ΔF/F_0_ during the 500 ms pre-cue epoch. To investigate the relationship between ΔF/F_0_ trajectories and reaction time, reaction times were split into thirds (short [0 – 350 ms], medium [350 - 900 ms] and long [>900 ms] and only FOVs with a sufficient number of trials per reaction time category were included in further analysis. The onset times of early enhanced neurons was calculated per trial for each FOV by employing a previously published onset detection algorithm using a slope sum function (SSF) (Zong et al., 2003) with the decision rule and window of the SSF adapted to calcium imaging data (threshold 10% of peak, SSF window 375 ms, smoothed with a Savitzky Golay filter across 27 frames with order 2). To reduce the influence of noisy individual traces biasing onset detection, each onset was determined as the median of 10,000 bootstrap samples. After calculating an onset for each trial, a kernel density estimate was calculated for the mean onset across trials. The area under this mean population kernel density estimate was calculated using numerical trapezoidal integration. The reliability index for each neuron was defined as the mean Pearson’s correlation coefficient across pairs of trials in a defined window from -500 to +500 ms peri-movement initiation. The signal-to-noise ratio was defined as the ratio of mean absolute peak ΔF/F_0_ change (1s pre-cue to 2s post-movement) and ΔF/F_0_ SD during the precue baseline. Time-to-half-maximum ΔF/F_0_ was calculated as the time from cue onset to 50% of the ΔF/F_peak_ trial-to-trial.

### *In vivo* electrophysiology

Whole-cell patch-clamp recordings targeted to layer 5B, 600–950 µm from the pial surface, were obtained from awake head restrained mice. Signals were acquired at 20 kHz using a Multiclamp 700B amplifier (Molecular Devices) and filtered at 10 kHz using PClamp 10 software in conjunction with a DigiData 1440 DAC interface (Molecular Devices). No bias current was injected during recordings and the membrane potential was not corrected for junction potential. Resting membrane potentials were recorded immediately after attaining the whole-cell configuration (break-in). Series resistances (Rs) ranged from 23.6 to 45.5 MΩ. Patch pipettes (5.5–7.5 MΩ) were filled with internal solution (285–295 mOsm) containing: 135 mM K-gluconate, 4 mM KCl, 10 mM HEPES, 10 mM sodium phosphocreatine, 2 mM MgATP, 2 mM Na_2_ATP, 0.5 mM Na_2_GTP, and 2 mg/ml biocytin (pH adjusted to 7.2 with KOH). External solution contained: 150 mM NaCl, 2.5 mM KCl, 10 mM HEPES, 1 mM CaCl_2_, and 1 mM MgCl_2_ (adjusted to pH 7.3 with NaOH). All electrophysiology recordings were analyzed using custom written scripts in MATLAB. Individual action potentials (APs) were detected with a wavelet-based algorithm (Nenadic and Burdick, 2005) and AP threshold was defined as the membrane potential (V_m_) at maximal d^2^V/dt^2^ up to 3 ms before AP peak and manually verified. For subthreshold V_m_ analysis APs were clipped by removing data points between -1 and +9 ms peri-AP threshold. Average AP firing frequencies were calculated by convolving spike times with a 50 ms Gaussian kernel. Significant changes in subthreshold V_m_ and AP firing frequency were defined by comparing bootstrapped 95% confidence intervals of mean movement-aligned V_m_ and AP frequency trajectories to zero (baseline epoch = 200 ms pre-cue; movement epoch = -100 to +100 ms peri-movement). Mean changes in V_m_ (ΔV_m_) were calculated by subtracting the mean V_m_ during baseline (1s epoch prior to cue) from the mean V_m_ during peri-movement epoch (-250 to +250 ms epoch when aligned to movement onset). All mean ΔV_m_ trajectories were decimated and median filtered with a 50 ms sliding window. Population mean ΔV_m_ trajectories were normalized to the largest absolute mean ΔV_m_ value in a 1.5 second peri-movement window. Peri-movement ΔV_m_ onsets were detected as the 10% rise-time of V_m_ trajectories when aligned to movement. To compare subthreshold V_m_ dynamics during hit and miss trials, cue-aligned periods of V_m_ were baseline subtracted and the area under the |ΔV_m_| trajectory from cue onset to median reward delivery was calculated via trapezoidal numerical integration with a 50 ms sample rate. We calculated the Pearson correlation coefficient between ΔV_m_ and Δfiring rate for all significantly modulated neurons.

### Immunohistochemistry

To morphologically identify neurons after recording, deeply anesthetized mice were transcardially perfused with 4 % paraformaldehyde. Mouse brains were post-fixed overnight and coronal sections (60 μm) of M1_FL_ were cut with a vibratome (Leica VT1000 S). For neuron location recovery, sections were incubated in streptavidin AlexaFluor-488 (1:1000, Molecular Probes) in 0.1 M phosphate buffered saline (PBS) containing 0.5 %Triton X-100, mounted (Vectashield, VectorLabs), imaged using a Zeiss LSM 510 Meta confocal microscope (20x objective) and referenced to the Franklin and Paxinos Mouse Brain Atlas (Paxinos & Franklin, 2008). To identify projection targets of individually recorded neuron (Schiemann et al., 2015), sections were further processed by heat-mediated antigen retrieval in 10 mM sodium citrate buffer (pH 6.0) for 3 hrs at 80°C. Sections were incubated in blocking solution (0.01 M PBS,10 % normal goat serum (v/v), 0.5 % Triton X-100 (v/v)) at 22 °C for 2 hrs and incubated overnight at 22 °C in a primary antibody mixture containing mouse monoclonal anti-Satb2 (1:200, Cat. No. ab51502, Abcam) and rat monoclonal anti-Ctip2 (1:1000, Cat. No. ab18465, Abcam) dissolved in carrier solution (0.01 M PBS, 1 % goat serum, 0.5 % Triton X-100). Slices were then incubated overnight at 22 °C in a secondary antibody mixture containing AlexaFluor-568 goat anti-mouse (1:750, Molecular Probes) and AlexaFluor-647 goat anti-rat (1:750, Molecular Probes) dissolved in carrier solution (0.01 M PBS, 1 % goat serum, 0.5 % Triton X-100), mounted and imaged using a Nikon A1R FLIM confocal microscope (Nikon, Europe). Images were analyzed offline using Fiji.

### Optogenetic experiments

For optogenetic activation of MTh_DN/IPN_ neurons or axon terminals in M1_FL_, 250 nl of AAV2/1-CAG-GhR2-GFP (4.7×10^11^ GC/ml, Addgene 20071; control virus: AAV2-CAG-mCherry (5.2×10^11^ GC/ml)) was injected unilaterally into contralateral MTh_DN/IPN_ (AP: -1.1, ML: 1.0, DV: -3.4 mm). For direct MTh_DN/IPN_ stimulation, an optic fiber (200 μm diameter, 0.39 NA; Thorlabs) was implanted ∼300 µm dorsal to the viral injection site and trains of pulsed 473 nm light (8 mW, 16.6 Hz pulse frequency, 33.3% duty cycle) were delivered using a solid-state laser (DPSS, Civillaser, China) and shutter (LS3S2T1, Uniblitz) controlled by an Arduino control system. For direct simulation of MTh_DN/IPN_ axon terminals, tapered optic fibers (Optogenix, Italy) were implanted to a depth of 1 mm at the center of M1_FL_ (AP: 0.6, ML: 1.6, DV: -1.0 mm) and 12 mW, 473 nm light was delivered with parameters as described above. Prior to optogenetic stimulation experiments, mice were trained to expert level performance and habituated to light emanating from an uncoupled optic fiber and the sound of shutter activation. During recording sessions, mice were exposed to 3 different trial types: (1) cue and shutter; (2) laser and shutter; and (3) shutter only. Trials were presented with the following pattern: 1, 1, 3, 1, 1, 2,… repeating for 30 minutes. For the majority of mice, direct MTh_DN/IPN_ stimulation was followed by MTh_DN/IPN_ neuron axon terminal stimulation in M1_FL_ on the following day. In some experiments, whole-cell patch-clamp recordings (as described above) were performed in combination with direct MTh_DN/IPN_ stimulation. To investigate behavioral context, mice which had previously experienced MTh_DN/IPN_ stimulation were head restrained above a 3D printed baseplate (Wanhao i3 Duplicator) without support/movable levers or reward spout and habituated to the altered behavioral context for 2 sessions, interleaved with normal training to ensure that the cued goal-directed motor behavior was not extinguished. To compare effects of MTh_DN/IPN_ stimulation in the learned and altered behavioral contexts, mice first underwent a 15 minute optogenetic stimulation protocol in the learned context, then returned to their home cage for 5 mins before being exposed to a 15 minute optogenetic stimulation protocol in the altered behavioral context. For histological confirmation of the injection site and optic fiber placement, mice were perfused and post-fixed for 2 additional days, before tissue slices were collected and imaged as described above. The center of the optic fiber (COF) was defined as the most ventral extent of the optic fiber tract across all slices from each brain as measured from the pial surface. Where tracts of equal depth were present, the coronal section containing the largest diameter tract tip was identified as the COF. The expression of ChR2-Venus was coarsely defined by first referencing three coronal slices (120 µm spacing) centered on the COF to the Franklin & Paxinos Mouse Brain Atlas (Paxinos & Franklin, 2008) before manually evaluating the proportion of each of the principle motor thalamic nuclei (AM, anteromedial; VL, ventrolateral; VPM, ventral posteromedial nucleus; VPL, ventral posteromedial; VM, ventromedial) containing fluorescence, and categorizing three levels based on expression covering 0%, 0-50% and 50-100% of each nucleus. Proportions of push-like movements in cue- and laser-trials were calculated by correcting for the behavioral error rate, i.e. subtracting the proportion of pushes observed in shutter only trials (3) to obtain a lower bound for induced movement proportion. ΔV_m_ trajectories for both cue-evoked and optogenetic stimulation-evoked movement trials were calculated as described above, and trial-by-trial ΔV_m_ changes were based on comparing the 200ms pre-laser or pre-cue epoch with the 200 ms perimovement epoch within each trial. We calculated the Pearson correlation coefficient between cue- and ChR2-evoked ΔV_m_ in hit trials.

### Forelimb kinematic tracking

Behavior from all experimental and habituation days was recorded at 300 frames per second using a high-speed camera (Pharmacological experiments: Genie HM640, Dalsa; optogenetic experiments: Mako U U-029, Allied Vision) and acquired with Streampix 7 (Norpix), synced using a TTL output from the DigiData 1440 DAC interface. Forepaw and wrist positions during pharmacological inactivation experiments were calculated by tracking forepaw markers using a custom written tracking script in Blender (2.79b, Blender Foundation). Contour plots of paw positions densities were calculated during 2 s epochs prior to cue presentation by sorting the paw positions by distance from the mean and computing 20 increasingly inclusive convex hulls around 5% portions of the data to define each contour level. Directional tracking of forelimb movement in the learned/altered behavioral context was performed using Deep Lab Cut, a markerless video tracking toolbox (Mathis et al., 2018). Paw trajectories were plotted for the 100 ms post movement onset epoch in the learned behavioral context (LBC), and for the altered behavioral context (ABC) we plotted trajectories in the epoch 100 ms after the LBC median reaction time, due to a lack of movement in the majority of ABC trials. Push-like movements were defined as trials with an initial paw trajectory vector between 100° and 210°. To measure gross forelimb movement, we defined a region-of-interest (ROI) covering the contralateral (left) forelimb and calculated the motion index (MI) for each successive frame *f* as 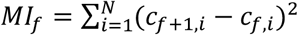, where c_*f,i*_ is the grayscale level of the pixel i of the ROI, N pixels per ROI^32^. Movement trials were defined by calculating the MI>*θ* within 500 ms of cue/laser onset, with the threshold *θ* defined as two standard deviations above mean MI.

### Statistics

Data analysis was performed using custom-written scripts in MATLAB 2019a and code will be made available on request. Data are reported as mean ± 95% bootstrapped confidence interval, 10,000 bootstrap samples, unless otherwise indicated. Where multiple measurements were made from a single animal, suitable weights were used to evaluate summary population statistics and to obtain unbiased bootstrap samples. Statistical comparisons using the significance tests stated in the main text were made in MATLAB 2019a, and statistical significance was considered when P<0.05 unless otherwise stated. Data were tested for normality with the Shapiro–Wilk test, and parametric/non-parametric tests were used as appropriate and as detailed in the text.

