## Supplemental Figures S1-6 for "Cerebellar-recipient motor thalamus drives behavioral context-specific movement initiation"

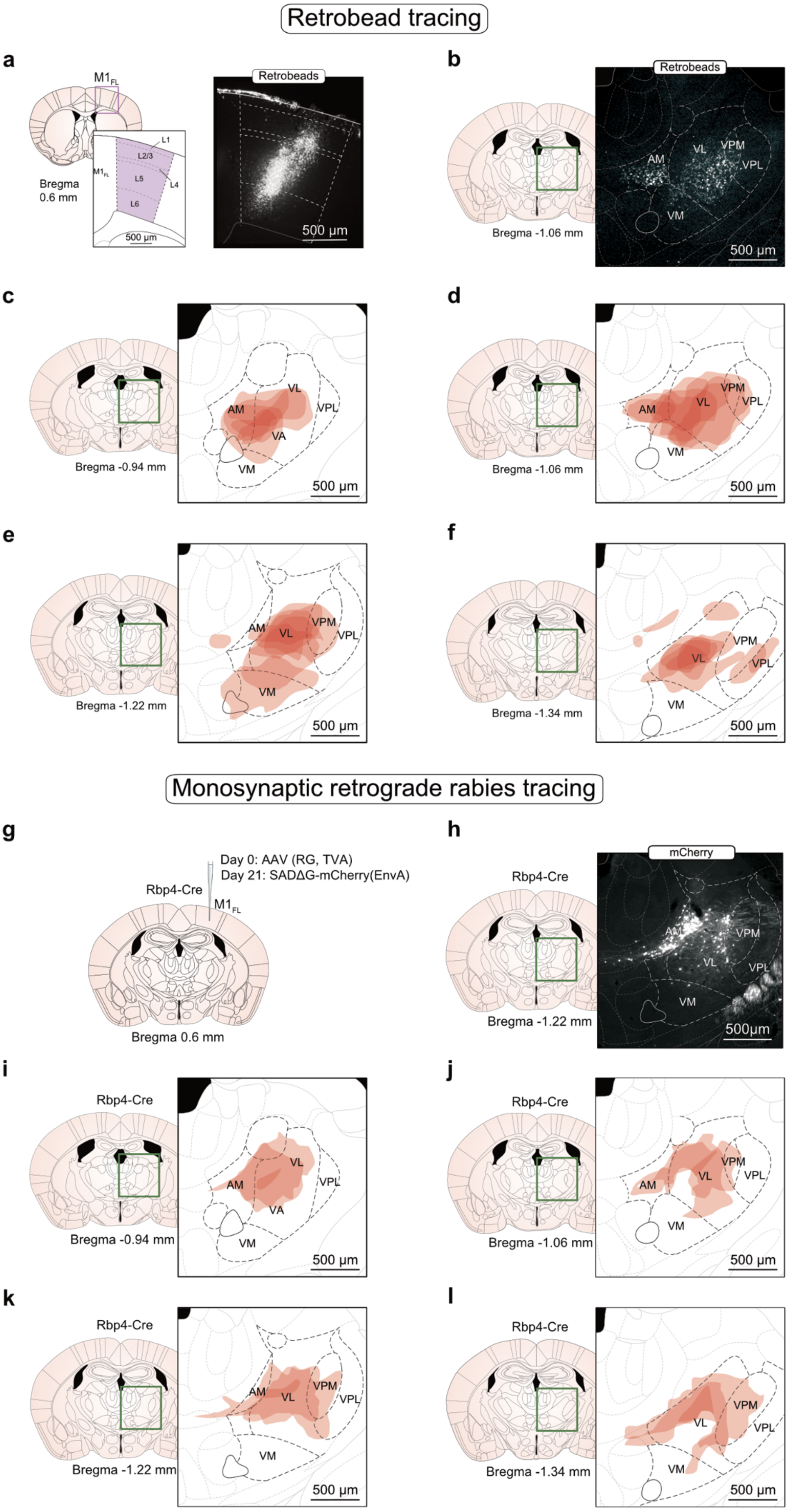
Retrobead tracing
a
M1FL
L1
L2/3
L5
L4
L6
M1FL
Bregma
0.6 mm
500 μm
Retrobeads
500 μm
b
Retrobeads
AM
VL
VPM
VPL
VM
Bregma -1.06 mm
500 μm
c
AM
VL
VPL
VA
VM
Bregma -0.94 mm
500 μm
d
AM
VL
VPM
VPL
VM
Bregma -1.06 mm
500 μm
e
AM
VL
VPM
VPL
VM
Bregma -1.22 mm
500 μm
f
VL
VPM
VPL
VM
Bregma -1.34 mm
500 μm
Monosynaptic retrograde rabies tracing
g
Day 0: AAV (RG, TVA)
Day 21: SADΔG-mCherry(EnvA)
Rbp4-Cre
M1FL
Bregma 0.6 mm
h
mCherry
Rbp4-Cre
AM
VPM
VL
VPL
VM
Bregma -1.22 mm
500μm
i
Rbp4-Cre
AM
VL
VPL
VA
VM
Bregma -0.94 mm
500 μm
j
Rbp4-Cre
AM
VPM
VL
VPL
VM
Bregma -1.06 mm
500 μm
k
Rbp4-Cre
AM
VL
VPM
VPL
VM
Bregma -1.22 mm
500 μm
l
Rbp4-Cre
VPM
VL
VPL
VM
Bregma -1.34 mm
500 μm

**Figure S1. Retrograde mapping of $M1_{FL}$-projecting motor thalamic neurons.**

(**a**) *Left*: Coronal mouse brain schematic depicting $M1_{FL}$ laminae. *Right*: representative coronal brain slice depicting retrobead injection sites in $M1_{FL}$ demonstrating coverage from L2/3-L6. **(b)** *Left:* Coronal mouse brain schematic, green square depicts the region of thalamus shown in the expanded view (*right). Right:* Representative coronal mouse brain slice displaying $M1_{FL}$-projecting motor thalamic neurons. **(c-f)** Distribution of $M1_{FL}$-projecting motor thalamic neurons labelled by fluorescent retrobeads (red, N = 5) for coronal positions (b) -0.94, (c) -1.06, (d) -1.22, (e) -1.34 mm relative to bregma. Green squares depict the region of thalamus shown in the expanded view (*right*). Motor thalamic nuclei: AM, anteromedial; VL, ventrolateral; VA, ventroanterior; VPM, ventral posteromedial nucleus; VPL, ventral posteromedial; VM, ventromedial. **(g)** Monosynaptic retrograde rabies tracing strategy: injection of AAV2/1-CAG-FLEX-mTagBFP2-2A-TVA & SADΔG-mCherry(EnvA) into forelimb motor cortex ($M1_{FL}$) of a layer 5-specific Rbp4-Cre mouse. **(h)** *Left:* Coronal mouse brain schematic, green square depicts the region of thalamus shown in the expanded view (*right). Right:* Representative coronal mouse brain slice displaying $M1_{FL}$-projecting motor thalamic neurons. **(i-l)** Distribution of $M1_{FL}$-projecting motor thalamic neurons labelled by monosynaptic retrograde rabies tracing (red, N = 3) for coronal positions (b) -0.94, (c) -1.06, (d) -1.22, (e) -1.34 mm relative to bregma. Green squares depict the region of thalamus shown in the expanded view (right). Motor thalamic nuclei: AM, anteromedial; VL, ventrolateral; VA, ventroanterior; VPM, ventral posteromedial nucleus; VPL, ventral posteromedial; VM, ventromedial.

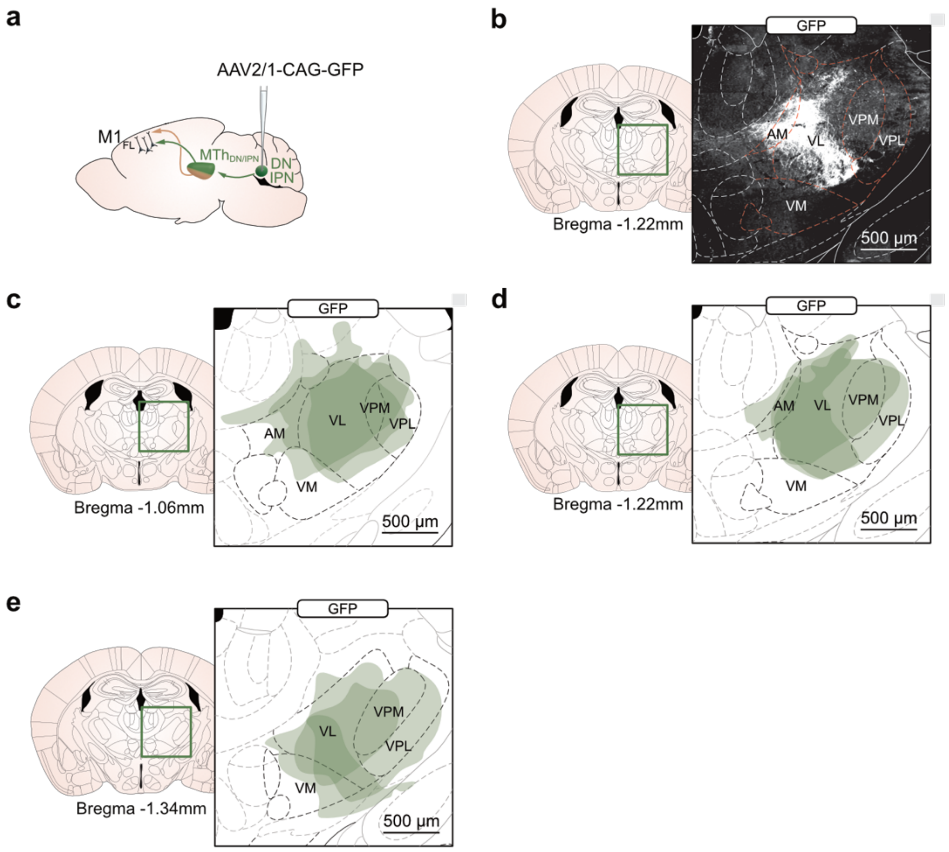


**Figure S2. Axon terminal fields of dentate / interpositus nucleus neurons in motor thalamus.**

(**a**) Sagittal mouse brain schematic illustrating viral-based anterograde mapping of dentate nucleus (DN) and interpositus nucleus (IPN) projections to motor thalamus. **(b)** *Left:* Coronal mouse brain schematic, green square depicts the region of thalamus shown in the expanded view (*right). Right:* Representative coronal mouse brain slice displaying anterograde mapping of dentate nucleus (DN) and interpositus nucleus (IPN) axonal arborisation in motor thalamic nuclei. **(c-e)** Distribution of labelled dentate nucleus and interpositus nucleus axon terminals in motor thalamus (N = 3 mice) for coronal positions (b) -1.06, (c) -1.22, (d) -1.34 mm relative to bregma. Green squares depict the region of thalamus shown in the expanded view (right). Motor thalamic nuclei: AM, anteromedial; VL, ventrolateral; VA, ventroanterior; VPM, ventral posteromedial nucleus; VPL, ventral posteromedial; VM, ventromedial.

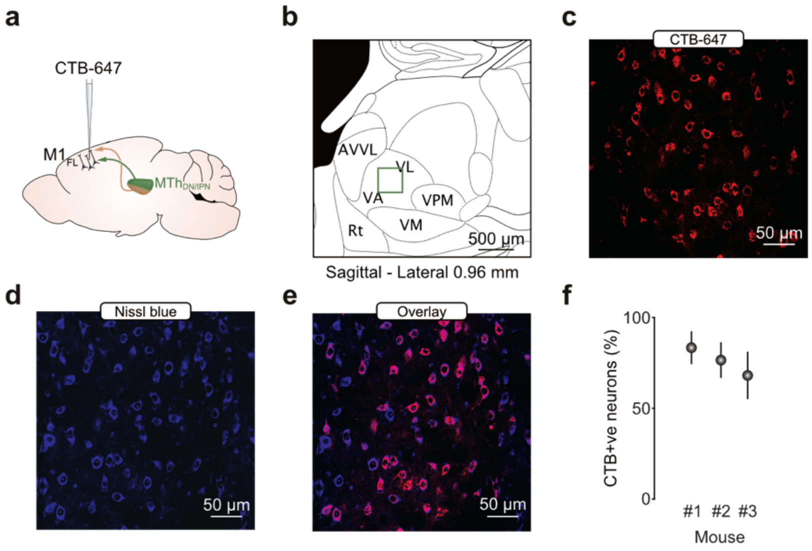


**Figure S3. Quantification of M1$_{FL}$-projecting neurons in ventroanterior / ventrolateral (VA/VL) region of motor thalamus.**

**(a)** Sagittal brain slice schematic illustrating retrograde cholera toxin b-Alexa647 (CTB-Alexa647) mapping of M1$_{FL}$-projecting neurons in ventroanterior / ventrolateral (VA/VL) region of motor thalamus. **(b)** Sagittal mouse brain atlas overlay of motor thalamic nuclei. Green square depicts the region of thalamus shown in the expanded view in c-e. **(c-e)** Motor thalamic neurons labelled with CTB-Alexa647 (c), Nissl blue counter stain (d) and overlay (e). **(f)** Proportion of M1$_{FL}$-projecting motor thalamic neurons in VA/VL (N = 3 mice, 4-6 slices per mouse, mean ± bootstrapped 95% CI).

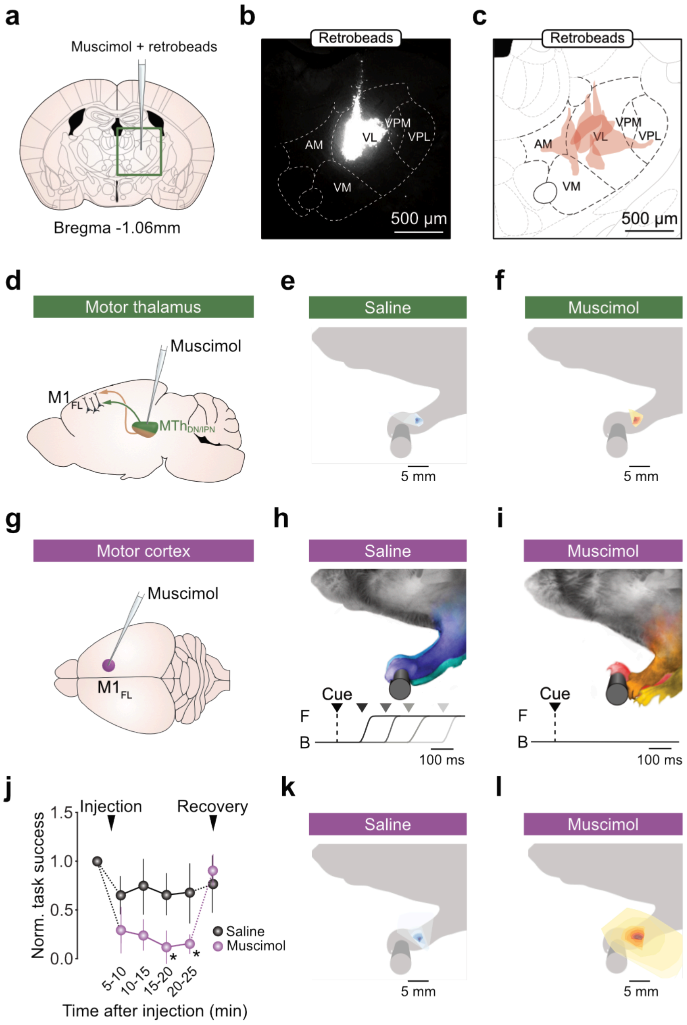


**Figure S4. Muscimol inactivation of primary motor cortex and dentate / interpositus nucleus-recipient area of motor thalamus.**

**(a)** Coronal brain schematic illustrating motor thalamus inactivation strategy. Green box identifies expanded view shown in (b-c). **(b)** Coronal brain slice displaying representative muscimol injection site within motor thalamus. **(c)** Distribution of muscimol injection sites (N = 5 out of 10 sites recovered). **(d)** Dentate / interpositus nucleus-recipient motor thalamus inactivation strategy. **(e-f)** Contour plot of baseline paw positions after (e) saline or (f) muscimol injection into $MTh_{DN/IPN}$. Each colored region of the contour plot contains 5% of the data. **(g)** Top-down view of a mouse brain illustrating $M1_{FL}$ muscimol inactivation strategy. **(h-i)** *Top:* Superimposed images of forelimb position at cue onset after (h) saline or (i) muscimol injection into $M1_{FL}$. *Bottom*: Example lever trajectories from 4 different trials (black to light grey) following injection of (h) saline or (i) muscimol into $M1_{FL}$. B, back lever position; F, front lever position; black triangle and dashed line represent cue presentation; grey triangles indicate reaction times. **(j)** Normalised task success as a function of time after muscimol injection. Colored symbols represent changes in population means ± 95% CI after saline (black, N = 5) or muscimol (purple, N = 5) injection (* $P < 0.05$, 2-way repeated measures ANOVA, Bonferroni-Holm correction for multiple comparisons). **(k-l)** Contour plot of baseline paw positions after (k) saline or (l) muscimol injection into $M1_{FL}$. Each coloured region of the contour plot contains 5% of the data.

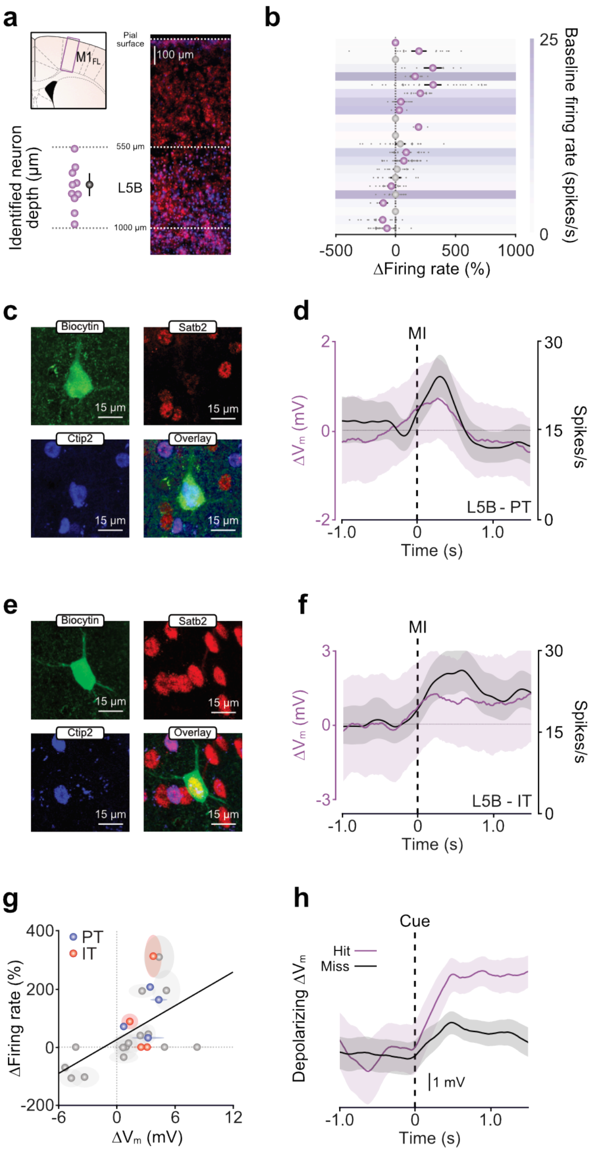


**Figure S5. Membrane potential dynamics and projection class identity of individually recorded L5B neurons.**

**(a)** *Top left*: Coronal brain slice schematic showing M1$_{FL}$. The purple rectangle depicts the expanded view (*right*). *Right:* Distribution of PT-type (blue, Ctip2 staining) and IT-type (red, Satb2 staining) projection neurons in L5B of M1$_{FL}$. *Bottom left:* depth of recovered L5B neurons as measured perpendicularly from the pial surface (n = 11/23 neurons identified, black symbol represents mean ± 95% CI). **(b)** Average change in L5B projection neuron peri-movement firing rate ± 95% CI. Grey dots represent individual trials, purple symbols represent significant changes in firing rate, white symbols represent non-significant changes, defined by comparing 95% bootstrapped confidence intervals. **(c)** Biocytin staining (green) and post-hoc immunohistochemical staining for Satb2 (red) and Ctip2 (blue) confirmed the PT-type projection class identity of an individually recorded L5B pyramidal neuron. **(d)** Mean subthreshold V$_m$ and firing rate of the L5B PT-type projection neuron depicted in (c). **(e)** Biocytin staining (green) and post-hoc immunohistochemical staining for Satb2 (red) and Ctip2 (blue) confirmed the IT-type projection class identity of an individually recorded L5B pyramidal neuron. **(f)** Mean subthreshold V$_m$ and firing rate of the L5B IT-type projection neuron depicted in (e). **(g)** Correlation between movement-related subthreshold ΔV$_m$ and firing rate changes. Blue/red symbols represent means ± 95% CI from individual PT-/IT-type neurons respectively, black line is a linear fit to the data. Grey symbols represent means ± 95% CI from neurons where projection class identity was not determined. **(h)** Example cue aligned hit-trial mean ΔVm (purple) vs miss-trial mean ΔVm (black) trajectories from a representative M1$_{FL}$ layer 5B neuron.

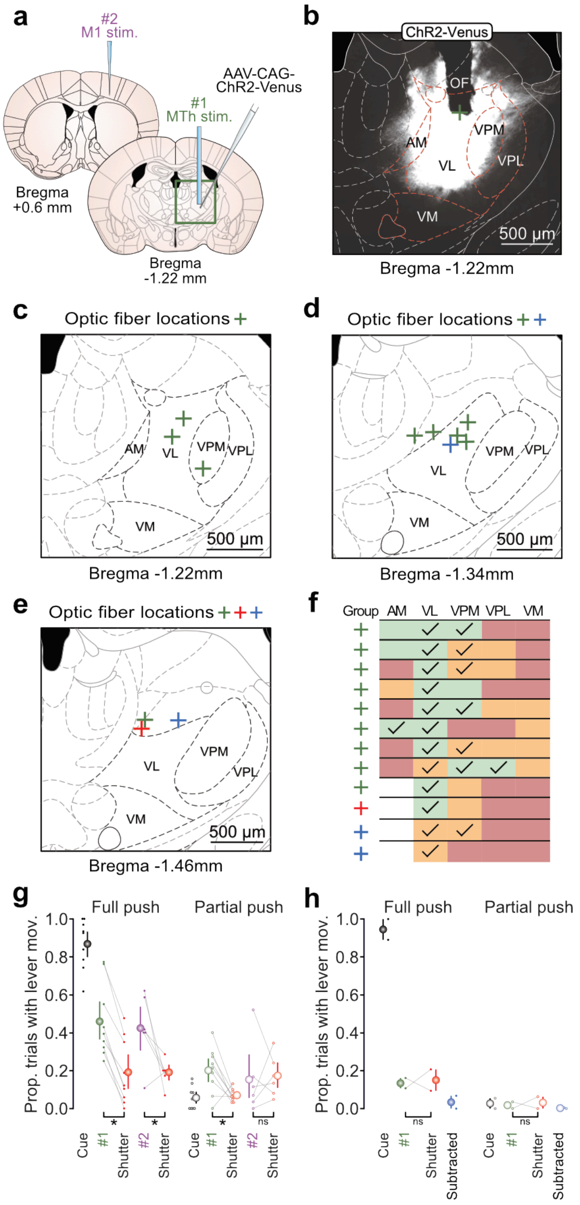


**Figure S6. Viral expression and selective optogenetic stimulation of dentate / interpositus nucleus-recipient region of motor thalamus.**

**(a)** Coronal brain slice schematics illustrating dual MTh optogenetic stimulation strategy: channelrhodopsin 2 (ChR2) expression was targeted to neurons in $MTh_{DN/IPN}$ and stimulated via an optic fiber placed directly above $MTh_{DN/IPN}$ (#1) or via a tapered optic fiber implanted directly into forelimb motor cortex ($M1_{FL}$) (#2). Green square overlay identifies the expanded view shown in (b). **(b)** Representative coronal section showing expression of ChR2-Venus and position of the optic fiber (OF), centre marked by green cross. **(c-e)** Anatomical locations of optic fiber placements in $MTh_{DN/IPN}$ (N = 12 mice, green, red and blue crosses indicate positive behavioral effects, no behavioral effects and control mice, respectively). **(f)** Quantification of viral expression in motor thalamic nuclei (green, nuclei with 50-100% coverage; orange, nuclei with 0-50% coverage; red, no expression; ticks represent nuclei that are within the cone of 470nm light emitted from the optic fiber during optogenetic stimulation). **(g)** Comparison of cue-evoked and ChR2-evoked task engagement represented as the proportion of trials that evoked either full (*left*) or partial (*right*) lever push movements. Black, cue + shutter trials; green, direct $MTh_{DN/IPN}$ stimulation + shutter trials; purple, direct stimulation of $MTh_{DN/IPN}$ axon terminals in $M1_{FL}$ + shutter trials; orange, shutter only trials. Colored dots represent data from individual mice, colored symbols represent mean ± 95% CI. (Cue, N = 9 mice; #1, N = 9 mice, #2, N = 6 mice. * $P$ < 0.05, ns = non-significant, two-sample $t$-test) **(h)** Comparison of cue-evoked and ChR2-evoked task engagement represented as the proportion of trials that evoked either full (*left*) or partial (*right*) lever push movements in control mice. Black, cue + shutter trials; green, direct $MTh_{DN/IPN}$ stimulation + shutter trials; orange, shutter only trials; blue, (#1 – shutter) trials. Colored dots represent data from individual mice, colored symbols represent mean ± 95% CI. (N = 2 mice, ns = non-significant, two-sample $t$-test).
